# Cell type-specific loops linked to RNA polymerase II elongation in human neural differentiation

**DOI:** 10.1101/2023.12.04.569731

**Authors:** Katelyn R. Titus, Zoltan Simandi, Harshini Chandrashekar, Dominik Paquet, Jennifer E. Phillips-Cremins

**Affiliations:** Department of Bioengineering, School of Engineering and Applied Science, University of Pennsylvania, Philadelphia, PA; Epigenetics Institute, Perelman School of Medicine, University of Pennsylvania; Institute for Stroke and Dementia Research, Ludwig Maximilians Universitat, Munich, Germany; Department of Genetics, Perelman School of Medicine, University of Pennsylvania

**Author notes:** Authors contributed equally to this work (Katelyn R. Titus, Zoltan Simandi, Harshini Chandrashekar).

## Abstract

DNA is folded into higher-order structures that shape and are shaped by genome function. The role for long-range loops in the establishment of new gene expression patterns during cell fate transitions remains poorly understood. Here, we investigate the link between cell-specific loops and RNA polymerase II (RNAPolII) during neural lineage commitment. We find thousands of loops decommissioned or gained *de novo* upon differentiation of human induced pluripotent stem cells (hiPSCs) to neural progenitors (NPCs) and post-mitotic neurons. During hiPSC-to-NPC and NPC-to-neuron transitions, genes changing from RNAPolII initiation to elongation are >4-fold more likely to anchor cell-specific loops than repressed genes. Elongated genes exhibit significant mRNA upregulation when connected in cell-specific promoter-enhancer loops but not invariant promoter-enhancer loops, promoter-promoter loops, or unlooped. Genes transitioning from repression to RNAPolII initiation exhibit slight mRNA increase independent of loop status. Our data link cell-specific loops and robust RNAPolII-mediated elongation during neural cell fate transitions.

**Highlights:**

- Thousands of loops are decommissioned and gained upon human iPSC differentiation to NPCs/neurons
- Genes transitioning from initiated-to-elongated exhibit robust mRNA upregulation when connected in cell type-specific promoter-enhancer loops
- Genes transitioning from repressed-to-initiated exhibit slight increases in mRNA levels independent of loop status
- Upon short-term RNAPolII degradation, loops formed by elongated genes are more severely disrupted than those anchoring initiated genes

## Introduction

Mammalian development requires the precise spatiotemporal regulation of gene expression in defined cell types. Non-coding cis-regulatory elements (CREs) known as enhancers regulate cell type-specific patterns of gene expression in metazoans^1–3^. CREs can be separated by kilobases (kb) up to megabases (Mb) from their target genes^4,5^, therefore understanding how chromatin loops are involved in governing enhancer-promoter communication is critical toward our knowledge of gene expression regulation.

There are three broad categories of long-range looping interactions: promoter-promoter (P-P) loops, enhancer-promoter (E-P) loops, and structural loops with no direct connection to promoters. In the case of structural loops, it is established that they are anchored by the architectural proteins CTCF and cohesin^6,7^. During interphase in steady-state mammalian cells, the cohesin complex extrudes DNA through its ring in an ATP-dependent manner to create transient “loops in the making”^8–11^. Cohesin-mediated loop extrusion stalls at CTCF-occupied motifs oriented toward each other in convergent orientation^10,12,13^. Extrusion stalling manifests empirically as focal ‘dot-like’ structures representing loop anchors in ensemble Hi-C heatmaps^14^. Short-term degradation of CTCF or the Rad21 subunit of cohesin ablates structural loops^15–17^. By contrast, only a subset of long-range E-P or P-P loops are anchored by CTCF and sensitive to its degradation^16^. The zinc finger-containing transcription factor Yin Yang 1 (YY1) and subunits of the Mediator complex have recently been linked to E-P loop maintenance, but results vary depending on the cell type and methodologies used for molecular perturbation^16–23^. Overall, the mechanisms governing E-P and P-P loops and how they are functionally linked to transcription remain important open questions.

Enhancers and promoters are enriched for genetic motifs encoding binding sites for transcription factors. Through the process of protein-protein interactions, motif-bound transcription factors at CREs and transcription start sites (TSSs) can recruit co-factors and RNAPolII^24^. Transcription establishment requires assembly of the pre-initiation complex at the gene promoter, Cdk7-mediated C-terminal phosphorylation of RNAPolII-serine 5 (Ser5P), and RNAPolII initiation^25–27^. RNAPolII transcribes promoter-proximal RNA until it pauses approximately 20-60 bp downstream of the TSS via the direct binding to negative elongation factor (NELF) and DRB-sensitivity-inducing factor (DSIF)^28,29^. Paused RNAPolII is released into productive elongation via the CDK9 catalytic domain of positive transcription elongation factor (P-TEFb), which phosphorylates serine 2 on the C-terminal domain of RNAPolII as well as NELF and DSIF^25–27^. RNAPolII-mediated enhancer RNAs have also been detected at CREs in mammalian cells^30^. Two recent studies demonstrated that short-term RNAPolII degradation can disrupt a subset of E-P loops in steady-state human cell lines^31,32^, suggesting a direct link between transcription and loop maintenance. However, the link between cell type-specific E-P and P-P loops and the establishment of new RNAPolII initiation and elongation patterns remains an important unanswered question.

Here, we set out to understand the relationship between loops and RNAPolII during the establishment of new transcriptional programs during changes in cell fate. We create genome-wide reference maps of long-range chromatin loops, gene expression, and RNAPolII occupancy during neural lineage commitment in the transitions from human induced pluripotent stem cells (hiPSCs) to neural progenitors (NPCs) and NPCs to post-mitotic neurons. We uncover a strong link among cell type-specific E-P loops gained *de novo* during differentiation, RNAPolII elongation, and robust increase of gene expression during human neural lineage commitment. We also demonstrate that loops anchored by elongated genes are particularly sensitive to short term RNAPolII degradation whereas loops anchoring initiated genes bound by CTCF are protected from RNAPolII perturbation. Our work sheds new light into the genome’s dynamic and context-dependent structure-function relationship by linking chromatin loops to transcription elongation during human neural lineage commitment.

## Results

### Rewiring of chromatin loops in a human iPSC model of early neural lineage commitment

We set out to conduct a genome-wide analysis of long-range chromatin loops in an *in vitro* human neurodevelopmental model consisting of three cellular stages: (1) undifferentiated human induced pluripotent stem cells (hiPSCs) (Day 0); (2) iPSC-derived neural progenitor cells (NPCs, DIV35) and (3) post-mitotic cortical neurons (neurons, DIV65) (**Figure 1A**). We implemented a well-established, multi-stage protocol for growth factor-mediated neuronal differentiation of iPSCs in monolayer tissue culture (**STAR Methods**)^33,34^. After confirming the absence of karyotypic abnormalities, we used immunofluorescence and microscopy to evaluate the efficiency of differentiation (**STAR Methods**). We observed that hiPSCs exhibit homogeneous expression of pluripotency markers SSEA4 and OCT3/4 (also known as POU5F1) (**Figure 1B**). We further observed that NPCs homogeneously express forebrain progenitor markers FOXG1 and NESTIN and that post-mitotic neurons showed homogenous expression of the pan-neural marker MAP2 and layer V-specific marker CTIP2 (**Figure 1B**). Thus, our hiPSC, NPCs, and neurons homogeneously display the expected morphology and protein markers characteristic of the stage of neural lineage commitment.

**Figure 1.**
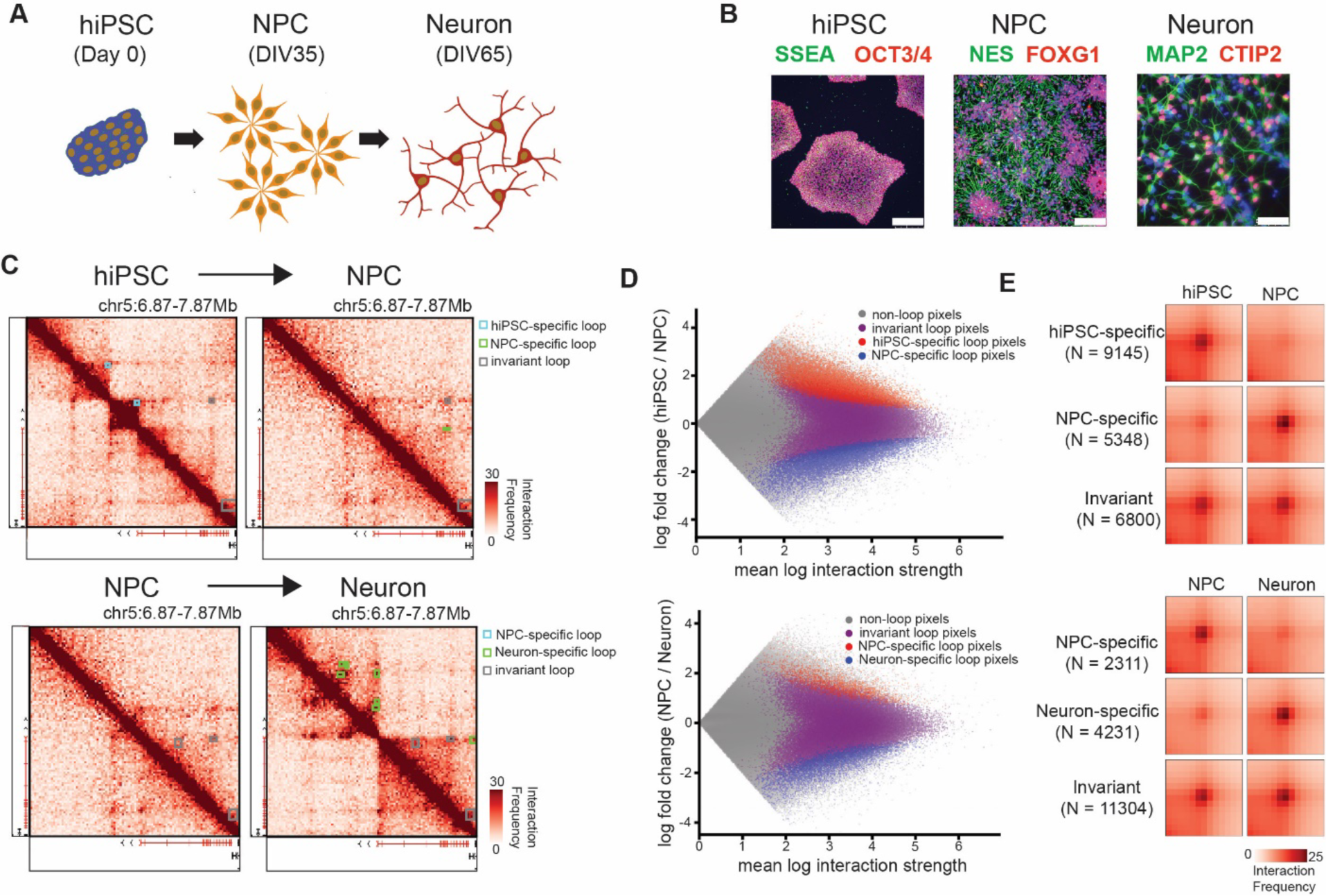
Differentiation of human iPSCs into neural progenitor cells and neurons leads to genome-wide rewiring of long-range chromatin loops. **(A)** A cartoon schematic of the differentiation from induced pluripotent stem cells (hiPSCs) to neural progenitor cell (NPCs) and neurons. DIV represents the days of in vitro differentiation at which the samples were collected. (**B)** Immunostaining of hiPSCs, NPCs, and neurons for cell type-specific markers. Scale bar, 250 µm. (**C**) Hi-C heatmaps at the *ADCY2* locus with annotated invariant and cell type-specific loops between hiPSC and NPCs (top row) and between NPCs and neurons (bottom row). *ADCY2* highlighted in red on the gene track below the heatmap. (**D**) MA plots of cell type-specific loops (**STAR Methods**) between hiPSC and NPCs (top row) and between NPCs and neurons (bottom row). Non-loops, grey. Invariant loops, purple. Cell type-specific loops, red and blue. **E**, Aggregate peak analysis (APA) of differential and invariant loops across hiPSC and NPC conditions (top three rows) and NPC and neuron conditions (bottom three rows).

We assayed higher-order chromatin folding genome-wide by creating Hi-C maps in hiPSCs, NPCs, and neurons. We acquired 396 (hiPSCs), 474 (NPCs), and 426 (neurons) million reads for two biological replicates per condition, achieving a read depth sufficient for high-resolution chromatin loop calling (**STAR Methods**). We merged replicates to create genome-wide ensemble interaction frequency maps for all chromosomes at 10 kilobase (kb) resolution. We employed statistical methods developed by our lab and others to identify dot-like structures representative of *bona fide* loops in ensemble Hi-C data^7,35–37^. Dots are characterized in Hi-C maps as punctate groups of adjacent pixels with significantly higher contact frequency compared to the surrounding local topologically associating domain (TAD) and subTAD structure. We identified 17,178 loops in hiPSCs, 12,827 loops in NPCs, and 14,752 loops in neurons (**Table S1, STAR Methods**). We verified that our loop calls were robust across a sweep of parameters ^37^ and verified by visual inspection of Hi-C heatmaps (**Figure 1C**).

We previously developed a statistical method, 3DeFDR, to identify loops with invariant or cell type-specific interaction frequency between two biological conditions^36^. To quantify changes in looping genome-wide, we used 3DeFDR to compare hiPSC to NPC conditions and NPC to neuron conditions (**Figure 1C-E**). We found 9,145 hiPSC-specific loops, 5,348 NPC-specific loops, and 6,800 invariant loops between the hiPSC to NPC transition (**Figure 1D-E top, Table S2**). We further identified 2,311 NPC-specific loops, 4,231 neuron-specific loops, and 11,304 invariant loops for the NPC-to-neuron transition (**Figure 1D-E bottom, Table S3**). Our cell type specific and invariant loop calls were confirmed by visual inspection of Hi-C heatmaps. At the *ADCY2* locus we found loops that were decommissioned (prototypical hiPSC-specific loops) and gained *de novo* (prototypical NPC-specific and neuron-specific loops) upon differentiation (**Figure 1C**). Aggregate Peak Analysis (APA) plots of average interaction frequency validated our invariant and cell type-specific loop calls across both hiPSC to NPC and NPC to neuron transitions (**Figure 1E, STAR Methods**). Together, these results demonstrate that differentiation of hiPSCs into NPCs and neurons results in substantial rewiring of loops genome-wide.

### Classifying genes into repressed, initiated, and elongated RNAPolII occupancy

We next set out to classify genes at isoform resolution by their signature patterns of RNAPolII occupancy. It is well established that repressed genes will exhibit minimal to no RNAPolII signal, initiated genes will exhibit strong peak-like signal at the TSS and minimal signal at the gene body, and elongated genes will exhibit an RNAPolII peak at the TSS as well as domain-like signal across the gene body ^27,38^. We generated RNAPolII ChIP-seq libraries in hiPSCs, NPCs, and neurons (**STAR Methods**). To stratify genes into repressed, initiated, and elongated categories, first we merged genes with the same TSSs and TESs into transcriptional units, thus circumventing redundancies due to shared TSSs. We next quantified (1) the maximum RNAPolII signal in a window from −2kb to the TSS, (2) the mean RNAPolII signal across the gene body normalized by gene length, and (3) the mean signal within a window at the gene’s 3’ end that is proportional to the size of the transcriptional unit normalized by base pairs (**Figures 2A-B**, detailed in the **STAR Methods**). We verified that the computational strategy is effective in parsing repressed genes devoid of RNAPolII signal, initiated genes with an RNAPolII peak at the promoter and depleted of signal along the gene body, and elongated genes with high RNAPolII signal at the TSS and in a domain-like occupancy pattern along the gene body (**Figure 2C, Table S4**). Thus, we have classified repressed, initiated, and elongated transcriptional units across all three cell types.

**Figure 2.**
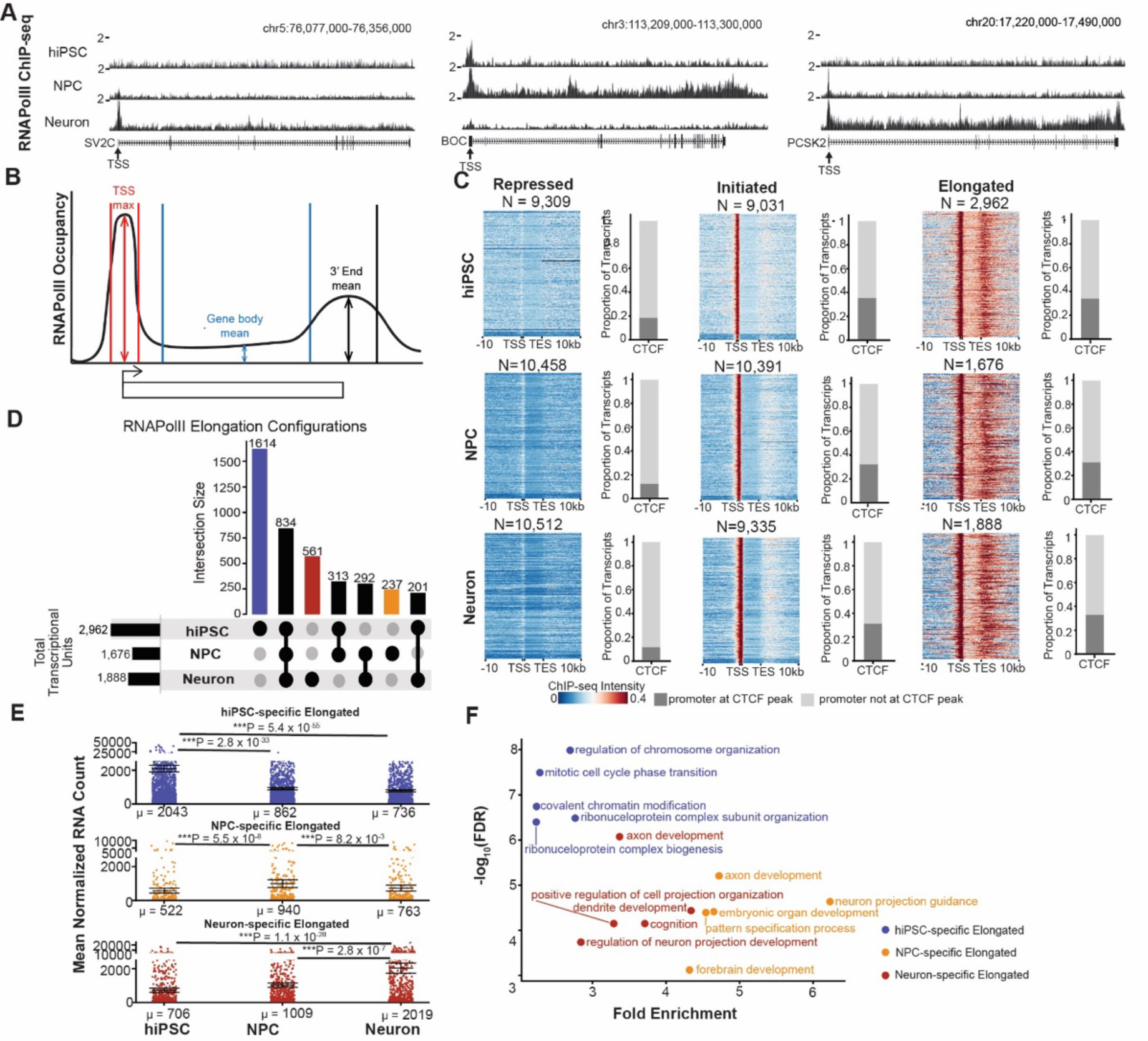
Categorization of transcriptional units by RNAPolII occupancy into repressed, initiated, and elongated. **(A)** Examples of transcriptional units exhibiting RNAPolII occupancy indicative of initiation in neurons (left), elongation in NPCs (middle), and elongation in neurons (right). **(B)** Schematic representing the windows used to categorize transcriptional units into repressed, initiated, and elongated by RNAPolII occupancy patterns (**STAR Methods**). (**C**) RNAPolII ChIP-seq heatmaps for transcriptional units categorized as repressed (left), initiated (middle), and elongated (right) in each cell type. Each row represents an individual transcriptional unit. Adjacent to the heatmaps are stacked bar plots illustrating the relative proportions of promoters co-localized with CTCF peaks. **(D**) Genes classified as elongated in hiPSC, NPC, and neurons. Horizontal bars, the number of unique transcriptional units exhibiting cell type-specific elongation. hiPSC-only (blue), NPC-only (orange), and neuron (red) elongated transcriptional units. (**E**) mRNA levels from RNA-seq for transcriptional units classified into blue, red, and orange in panel D. Points represent the mean normalized RNA count from three biological replicates. Horizontal lines represent the mean across all points. P-values, two-tailed Mann Whitney U (MWU) test with significance based on ɑ = 0.01. **(F)** Gene ontology analysis for transcriptional units classified into blue, red, and orange in panel D.

We next assessed CTCF occupancy patterns at the promoters of repressed, initiated, and elongated genes. Using CTCF ChIP-seq in hiPSC, NPCs, and neurons, we intersected the 2 kb segment upstream of all classified TSSs with CTCF peaks (**Figure 2C, Table S5**). Across the three cellular stages, we observed that less than 20% of promoters for repressed transcriptional units are occupied by CTCF. Promoters of initiated and elongated transcriptional units exhibit increases in CTCF occupancy compared to repressed genes. Our observations suggest that promoter occupancy of the architectural protein CTCF increases during the transition from repressed to initiated, but no substantial increase in genome-wide promoter occupancy occurs during the transition from initiation to elongation. Thus, it is unlikely that CTCF binding at the promoter alone could distinguish between initiated and elongated genes.

We sought to verify the elongated gene group by measuring their mRNA levels with bulk RNA-seq. We further stratified non-redundant, unique transcriptional units by those that were elongated only in hiPSCs, only in NPCs, or only in neurons (**Figure 2D)**. We found that hiPSC-specific elongated transcriptional units exhibited significantly higher mRNA levels in hiPSCs (µ = 2043; 95% confidence interval (CI): 1852 < μ_hiPSC_< 2235) compared to NPCs (µ = 862; 95% confidence interval (CI): 786 < μ_NPC_ < 937; Mann Whitney U (MWU), p = 2.8 x 10^−33^) and neurons (µ = 736; 95% confidence interval (CI): 670 < μ_neuron_< 802; MWU, p = 5.4 x 10^−55^). Similarly, we found that NPC-specific elongated transcriptional units had significantly higher expression in NPCs (µ = 940; 95% confidence interval (CI): 711 < μ_NPC_< 1169) compared to hiPSCs (µ = 522; 95% confidence interval (CI): 373 < μ_hiPSC_< 671; MWU, p = 5.5 x 10^−8^) and neurons (µ = 673; 95% confidence interval (CI): 493 < μ_neuron_< 852; MWU, p = 8.2 x 10^−3^). neuron-specific elongated transcriptional units exhibited significantly higher expression in neurons (µ = 2019; 95% confidence interval (CI): 1719 < μ_neuron_< 2319) compared to hiPSCs (µ = 706; 95% confidence interval (CI): 582 < μ_hiPSC_< 830; MWU, p = 1.1 x 10^−28^) and NPCs (µ = 1009; 95% confidence interval (CI): 884 < μ_NPC_< 1135; MWU, p = 2.8 x 10^−7^) (**Figure 2E**). All three groups of cell type-specific elongated genes showed the expected ontology for each cellular state (**Figure 2F)**, thus verifying our approach for stratifying elongated genes by RNAPolII occupancy.

### Dynamic RNAPolII occupancy during hiPSC-to-NPC and NPC-to-neuron cell fate transitions

Terminal differentiation of proliferating NPCs to post-mitotic neurons represents a unique cell fate transition in which chromatin is no longer subjected to the cell cycle. Focusing first on the NPC-to-neuron transition, we characterized genes into categories indicative of dynamic RNAPolII occupancy during cell fate transitions, including: (i) repressed (NPCs) to initiated (neurons), (ii) initiated (NPCs) to elongated (neurons), (iii) elongated (NPCs) to repressed (neurons), and (iv) constitutively repressed in both NPCs and neurons (**Figure 3A-E**). We confirmed that the transcriptional units constitutively repressed in the NPC-to-neuron transition displayed minimal RNAPolII occupancy (**Figure 3A**) and negligible mRNA levels (**Figure 3E**). Genes that transition from repressed-to-initiated during the NPC-to-neuron transition gained a RNAPolII peak at the promoter in neurons (**Figure 3B**).

**Figure 3.**
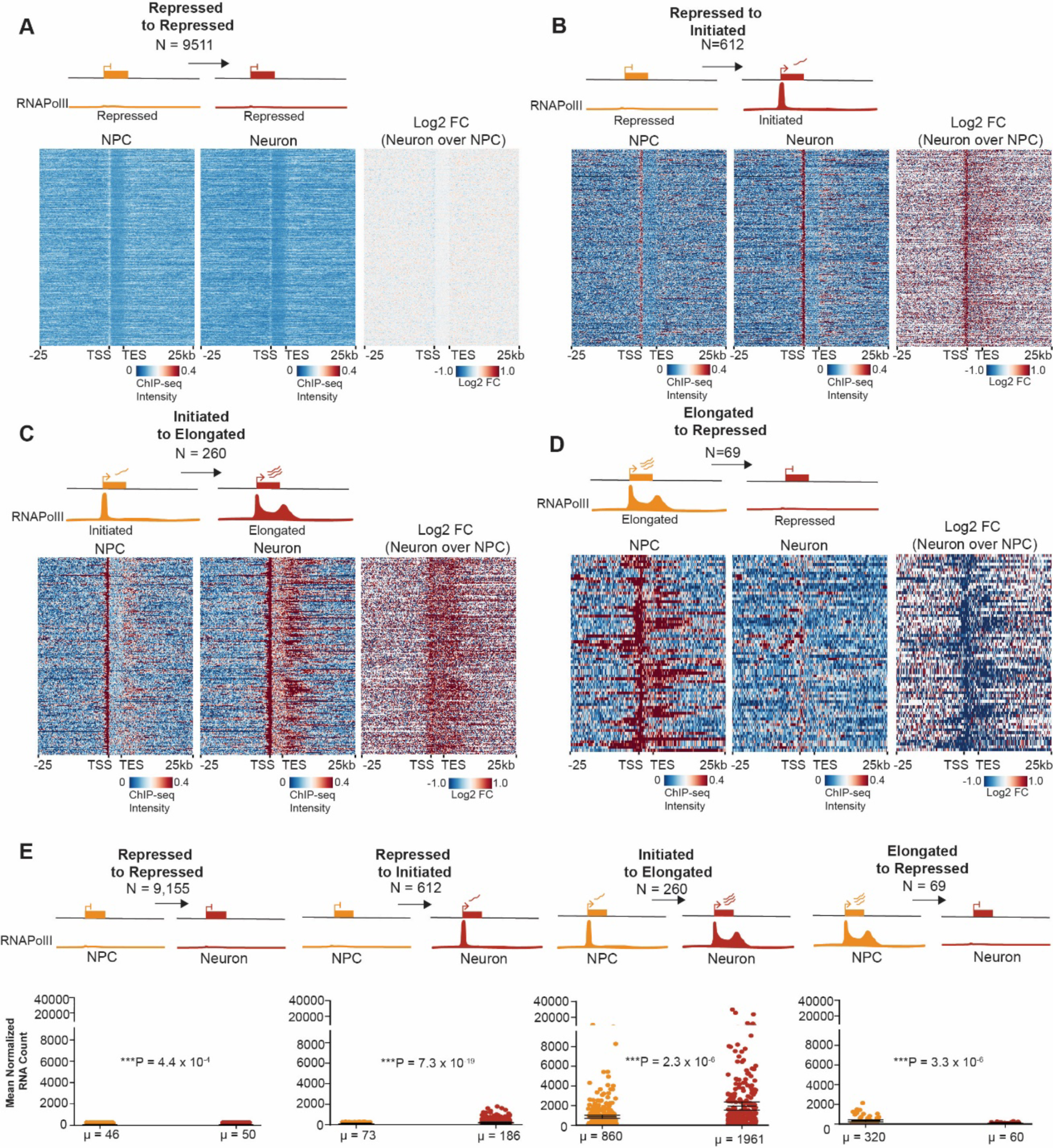
Dynamic RNAPolII occupancy patterns during the cell fate transition from human NPCs to post-mitotic neurons. (**A-D**) RNAPolII ChIP-seq heatmaps for transcriptional units categorized during the transition from iPSC-NPCs to post-mitotic neurons, including **(A)** repressed (NPCs) to repressed (neurons), **(B)** repressed (NPCs) to initiated (neurons), **(C)** initiated (NPCs) to elongated (neurons), and **(D)** elongated (NPCs) to repressed (neurons). Each row represents an individual transcriptional unit. (**E**) mRNA levels from RNA-seq for transcriptional units classified in A-D. Points represent the mean normalized RNA count from three biological replicates. Horizontal lines represent the mean across all points. P-values, two-tailed Mann Whitney U (MWU) test with significance based on ɑ = 0.01.

Moreover, genes that transition from initiated-to-elongated gained a pronounced domain-like pattern of RNAPolII signal along the gene body in neurons along with a significant upregulation in mRNA levels (**Figures 3C-E**). We also identified a group of genes displaying occupancy patterns consistent with RNAPolII decommissioning from elongation in NPCs to minimal RNAPolII occupancy in neurons, and these genes exhibit the expected downregulation in mRNA levels (**Figures 3D-E**). Such patterns were not restricted to the NPC-to-neuron transition, as the same RNAPolII occupancy and gene expression patterns for all four gene classes were similarly identified in the hiPSC-to-NPC transition (**Figure S1A-E**). Taken together, these data uncover gene classes with distinct dynamic RNAPolII occupancy patterns during hiPSC-to-NPC and NPC-to-neuron cell fate transitions.

### Genes transitioning from initiated to elongated during neural differentiation are strongly enriched for gained cell type-specific looping interactions

To understand the link between loops and RNAPolII occupancy, we stratified promoters of unique transcriptional units into those that are anchoring any NPC-specific (Class I, blue), Neuron-specific (Class 2, green), mixed (Class 3, yellow), cell type-invariant (Class 4, dark grey) chromatin loops and those not engaged in any looping interactions (Class 5, light grey) (**Figure 4A, Figure S2, STAR Methods**). At baseline, 29.5% of constitutively repressed transcriptional units connect in cell type-specific or invariant loops, whereas a striking 79.2% of initiated-to-elongated transcriptional units connect in cell type-specific or invariant loops during the NPC-to-neuron transition (**Figure 4B** first vs. third barplot, **Figure S2**). Transcriptional units transitioning from repressed-to-initiated exhibit equal probability of looping versus not looping to distal enhancers (**Figure 4B**, second barplot). By contrast, transcriptional units transitioning from initiated-to-elongated are >4-fold more likely to form cell type-specific loops versus repressed transcriptional units (**Figure 4B**, third barplot). Our data indicate that the majority of transcriptional units with RNAPolII occupancy patterns indicative of elongation anchor invariant or cell type-specific loops, whereas RNAPolII initiation occurs with similar likelihood in transcriptional units that loop or do not loop.

**Figure 4.**
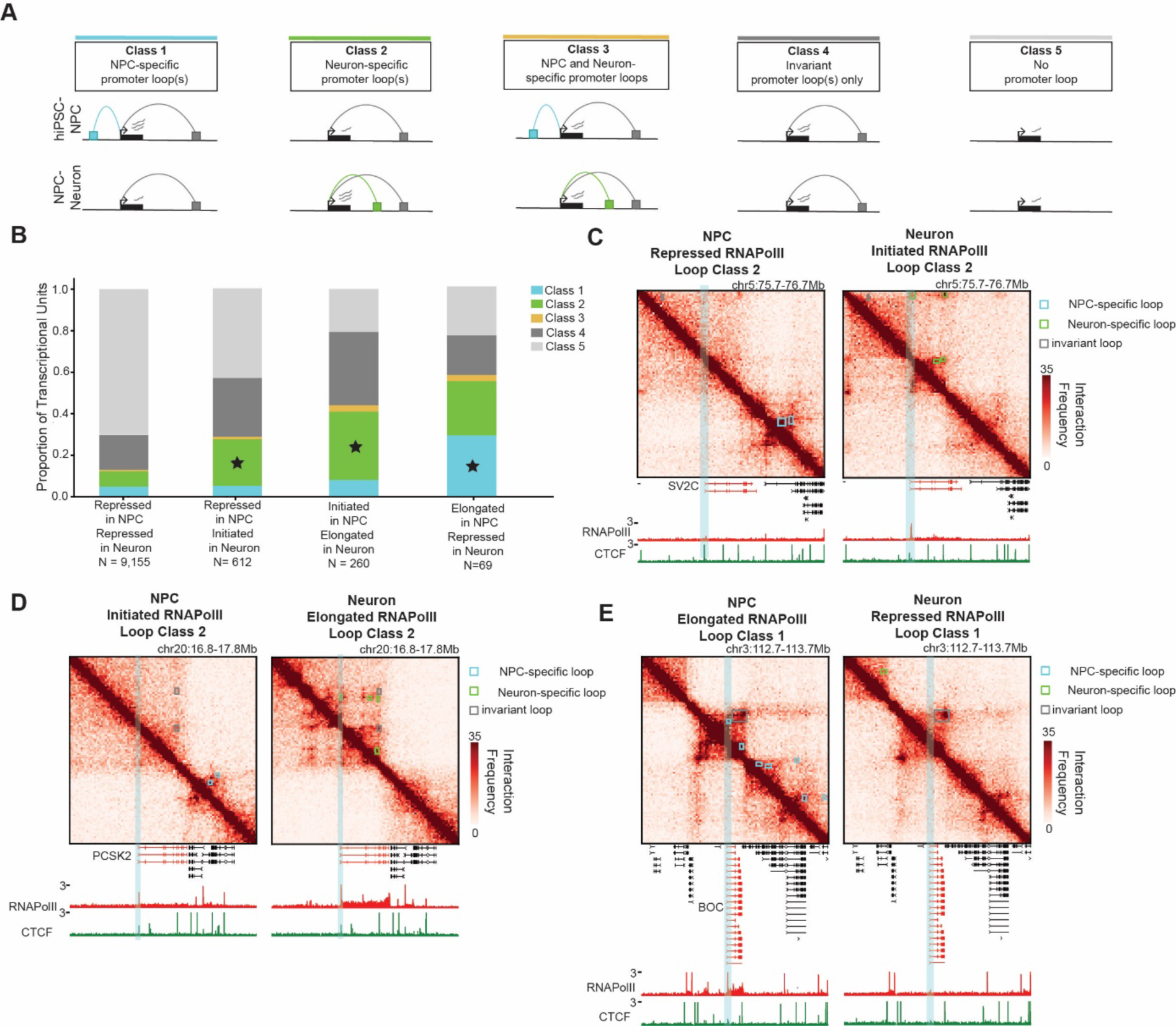
Genes transitioning from initiated-to-elongated are >4-fold more likely to form cell type-specific loops than repressed genes. **(A)** Schematic depicting promoters of unique transcriptional units classified as anchoring NPC-specific loops (Class 1, blue), neuron-specific loops (Class 2, green), mixed loops (Class 3, yellow), cell type-invariant loops (Class 4, dark grey), or not looping (Class 5, light grey). **(B)** Proportion of genes engaged in the five looping categories from panel 4A. Asterisk (*) indicates loop classifications of interest illustrated in panels 4C-E. **(C-E**) Hi-C heatmaps during NPC-to-neuron differentiation **(C)** the *SV2C* gene illustrating RNAPolII occupancy characteristic of the repressed-to-initiated cell fate transition, **(D)** the *PCSK2* gene illustrating RNAPolII occupancy characteristic of initiated-to-elongated genes, (**E)** the *BOC* gene illustrating RNAPolII occupancy characteristic of elongated-to-repressed transition. Tracks below Hi-C heatmaps show RNAPolII and CTCF ChIP-seq data.

A noteworthy observation is that both initiated and elongated genes are enriched for cell type-specific chromatin loops which arise *de novo* during neural differentiation. We observed that 22.3% of transcriptional units transitioning from repressed-to-initiated RNAPolII occupancy (**Figure 4B**, second barplot) and 33% of transcriptional units transitioning from initiated-to-elongated RNAPolII occupancy (**Figure 4B**, third barplot) gain neuron-specific loops that form *de novo* during the NPC-to-neuron transition (e.g. green Class 2 neuron-specific loops). By contrast, only 7.5% of transcriptional units constitutively repressed in the NPC-to-neuron transition engage in neuron-specific class 2 loops (**Figure 4B**, first barplot). Example neuron-specific loops gained in the NPC-to-neuron transition can be observed at the *SV2C* gene classified as transitioning from repressed to initiated (**Figure 4C**) and the *PCSK2* gene classified as transitioning from initiated to elongated (**Figure 4D**). We observed similar trends during the iPSC-to-NPC cell fate transition (**Figure S3A-E, Figure S4**). Our data suggest that transcriptional units are 3-4-fold more likely to engage in cell type-specific chromatin loops when they are initiated or elongated compared to repressed during hiPSC-to-NPC and NPC-to-neuron cell fate transitions.

### Genes transitioning from elongated to repressed during neural differentiation are strongly enriched for decommissioned cell type-specific looping interactions

We also examined the relationship between loops and genes transitioning from elongated-to-repressed during NPC-to-neuron differentiation. Transcriptional units that are elongated in NPCs and lose RNAPolII occupancy in neurons are strongly enriched in loops that are decommissioned in neurons (e.g. blue Class 1 NPC-specific loops) (**Figure 4B**). Specifically, we observe that 28.9% of decommissioned transcriptional units also lose loops, which is 6.5-fold higher than the 4.3% of repressed transcriptional units that engage in Class 1 NPC-specific loops (**Figure 4B**, fourth barplot, blue). Loop decommissioning is exemplified at the *BOC* gene classified with an elongated-to-repressed RNAPolII occupancy pattern (**Figure 4E**). Taken together, we observe that the establishment of new cell type-specific loops correlates with transitions into initiated and elongated RNAPolII and the decommissioning of loops correlates with the loss of RNAPolII occupancy during neural differentiation.

### Genes transitioning from repressed-to-initiated exhibit slight increases in mRNA levels independent of loop status

We next set out to distinguish between cell type-specific E-P and P-P loops for their relationship to RNAPolII and mRNA levels. We stratified the NPC-specific, neuron-specific, and invariant loops from **Figure 4A** and iPSC-specific, NPC-specific, and invariant loops from **Figure S3** into the subset connecting enhancers to promoters (E-P loops) (**Figure 5A-C, Figure S5, Figure S6, Figure S7A-C**) and those connecting promoters to other promoters (P-P loops) (**Figure 5D-F, Figure S5, Figure S6, Figure S7D-F**). We integrated E-P and P-P loops with genes classified by changes in RNAPolII occupancy and examined mRNA levels for both hiPSC-to-NPC (**Figures S6-S7**) and NPC-to-neuron (**Figures 5 and S5**) transitions (**STAR Methods**).

**Figure 5.**
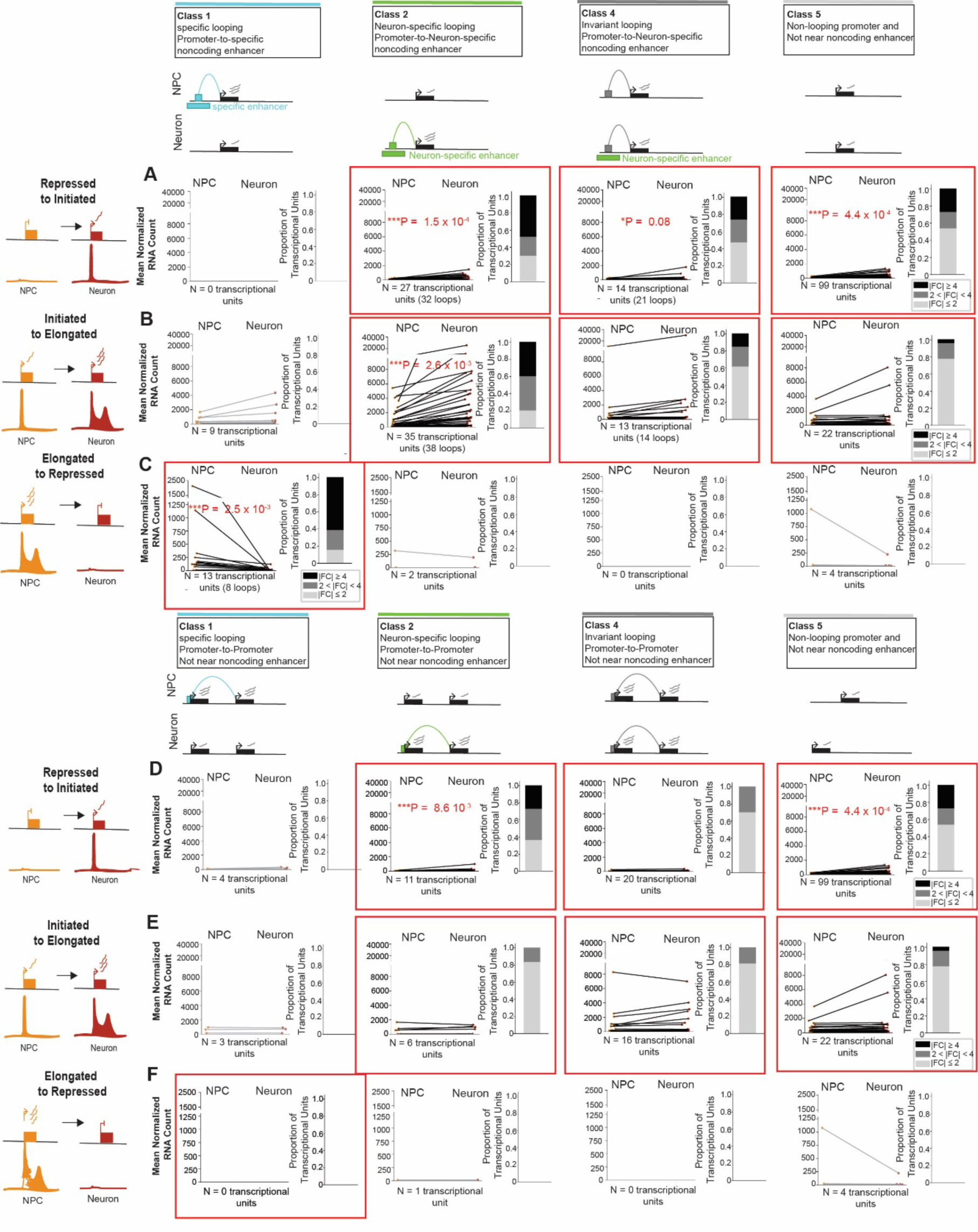
Upregulation of mRNA levels is highest for elongated genes engaged in de novo cell type-specific enhancer-promoter loops during NPC-to-neuron differentiation. (**A-F**) mRNA levels calculated from RNA-seq for transcriptional units stratified as (**A, D**) repressed to initiated, (**B, E**) initiated to elongated, and (**C, F**) elongated to repressed during NPC to neuron differentiation. (**A-C**) These three transcript classes are stratified into those that (column 1) anchor NPC-specific enhancer-promoter loops decommissioned in differentiation (Class 1, blue), (column 2) anchor neuron-specific enhancer-promoter loops gained de novo in differentiation (Class 2, green), (column 3) anchor cell type-invariant loops connecting promoters to neuron-specific promoters (Class 4, dark grey), and (column 4) do not loop and lack enhancers within a 80kb vicinity of the TSS (Class 5, light grey). (**D-F**) These three transcript classes are stratified into those that (column 1) anchor NPC-specific promoter-promoter loops decommissioned in differentiation (Class 1, blue), (column 2) anchor neuron-specific promoter-promoter loops gained de novo in differentiation (Class 2, green), (column 3) anchor cell type-invariant promoter-promoter loops (Class 4, dark grey), and (column 4) do not loop and lack enhancers within a 80kb vicinity of the TSS (Class 5, light grey). Each point is the mean normalized RNA count across three replicates. P-values are computed using a two-tailed Mann Whitney U test. Asterisk (*) reach significance under ɑ = 0.1 and triple asterisks (***) reach significance under ɑ = 0.01. Stacked bar plots represent the proportion of transcriptional units that exhibit absolute fold-change expression less than 2, greater than 2 and less than 4, and greater than 4 between NPC and neuron cellular states. Groups with less than 10 transcriptional units in panels A-C are greyed out due to the low number of transcriptional units. Red boxes highlight conditions with sample sizes sufficient to assess trends.

Several observations were revealed by our integrative analyses. First, for transcriptional units transitioning from repressed-to-initiated RNAPolII occupancy patterns, we observed a slight increase in mRNA levels whether the transcriptional units engaged in cell type-specific E-P loops, invariant E-P loops, or did not loop (**Figure 5A, Figure S7A**). We also observed slight increases in mRNA levels whether the transcriptional units engaged in E-P or P-P loops (**Figure 5D, Figure S7D**). Trends were similar in both the hiPSC-to-NPC and NPC-to-neuron transitions (**Figure 5, Figure S7**). Our data suggest that the recruitment of RNAPolII to the TSS in the repressed-to-initiated transition during neural differentiation can occur independent of looping status.

### Genes transitioning from initiated-to-elongated exhibit robust mRNA upregulation primarily when connected in cell type-specific enhancer-promoter loops

We next examined genes transitioning from initiated-to-elongated RNAPolII occupancy. We observed a striking upregulation of mRNA levels for those connected in cell type-specific E-P loops (**Figure 5B, Figure S7B**, second column). Similar patterns of mRNA upregulation did not occur for initiated-to-elongated genes when connected in P-P loops, invariant E-P loops, or not looping (**Figure 5B, Figure S7B**, fourth column; **Figure 5E, Figure S7E**, second column). Observations were similar in both the hiPSC-to-NPC and NPC-to-neuron transitions (**Figure 5, Figure S7**). Together these data indicate that although genes transitioning from repressed-to-elongated exhibit slight increases in mRNA levels independent of looping status, the transition from initiation-to-elongation correlates with significantly higher levels of steady state mRNA levels when the transcriptional units are engaged in cell type-specific E-P loops during neural differentiation.

### Decommissioning from elongation to repression can involve the breaking of cell type-specific E-P loops and can also occur when genes do not engage in loops

We also investigated genes transitioning from elongated-to-repressed RNAPolII occupancy. We observed downregulation of mRNA levels whether the transcriptional units engaged in decommissioned cell type-specific E-P loops or did not loop (**Figure 5C, Figure S7C**, first barplot). We did not find consistent decreases in mRNA levels when decommissioned genes engaged in invariant E-P loops or P-P loops (**Figure 5E, Figure S7E**). Similar trends were observed in the iPSC-to-NPC and NPC-to-neuron transitions (**Figure 5, Figure S7**). These results demonstrate that the decommissioning from productive elongation can involve the breaking of cell type-specific E-P loops and can also occur when the genes do not engage in loops.

### Short-term RNAPolII degradation disrupts loops anchored by elongated genes

Finally, we sought to functionally test our observation of a strong correlation among RNAPolII elongation, mRNA levels, and cell type-specific loops between enhancers and promoters of elongated genes. We re-analyzed published Micro-C data from the DLD-1 cell line in which an auxin-inducible degron was used to degrade RNAPolII^32^. Using published RNAPolII ChIP-seq^39^ and CTCF CUT&Tag^32^ data in the same DLD-1 cells, we stratified transcriptional units into those with or without CTCF promoter occupancy and exhibiting repressed, initiated, and elongated RNAPolII occupancy patterns (**Figure 6A-C**). We found that similarly to our NPC and neuron model system, 6% of repressed, 46% of initiated, and 64% of elongated transcriptional units genome-wide have CTCF bound at the promoter in DLD-1 cells (**Figure S8**). We also called loops (N = 24,358) genome-wide in Micro-C data in control DLD-1 cells without auxin (**STAR Methods**). Focusing on transcriptional units at the base of loops, we found that 27% of repressed transcriptional units, 68% of initiated transcriptional units, and 76% of elongated transcriptional units have at least one CTCF peak at the promoter (**Figure S8, Figure 6A-C**). Transcriptional units with CTCF at the promoter are more likely to anchor loops than those without CTCF (**Figure S8**). Loops anchoring repressed and initiated transcriptional units exhibit negligible to slight changes in interaction frequency upon lost RNAPolII occupancy, respectively (**Figure 6A-B**). By contrast, we observed a striking decrease in interaction frequency of loops anchoring elongated genes upon degradation of RNAPolII occupancy (**Figure 6A-C**). Elongated transcriptional units devoid of CTCF are particularly susceptible to loop disruption upon RNAPolII degradation (**Figure 6C**). These results further reinforce that RNAPolII is necessary for the maintenance of full loop interaction frequency, particularly for elongated genes that do not have CTCF bound at the promoter.

**Figure 6.**
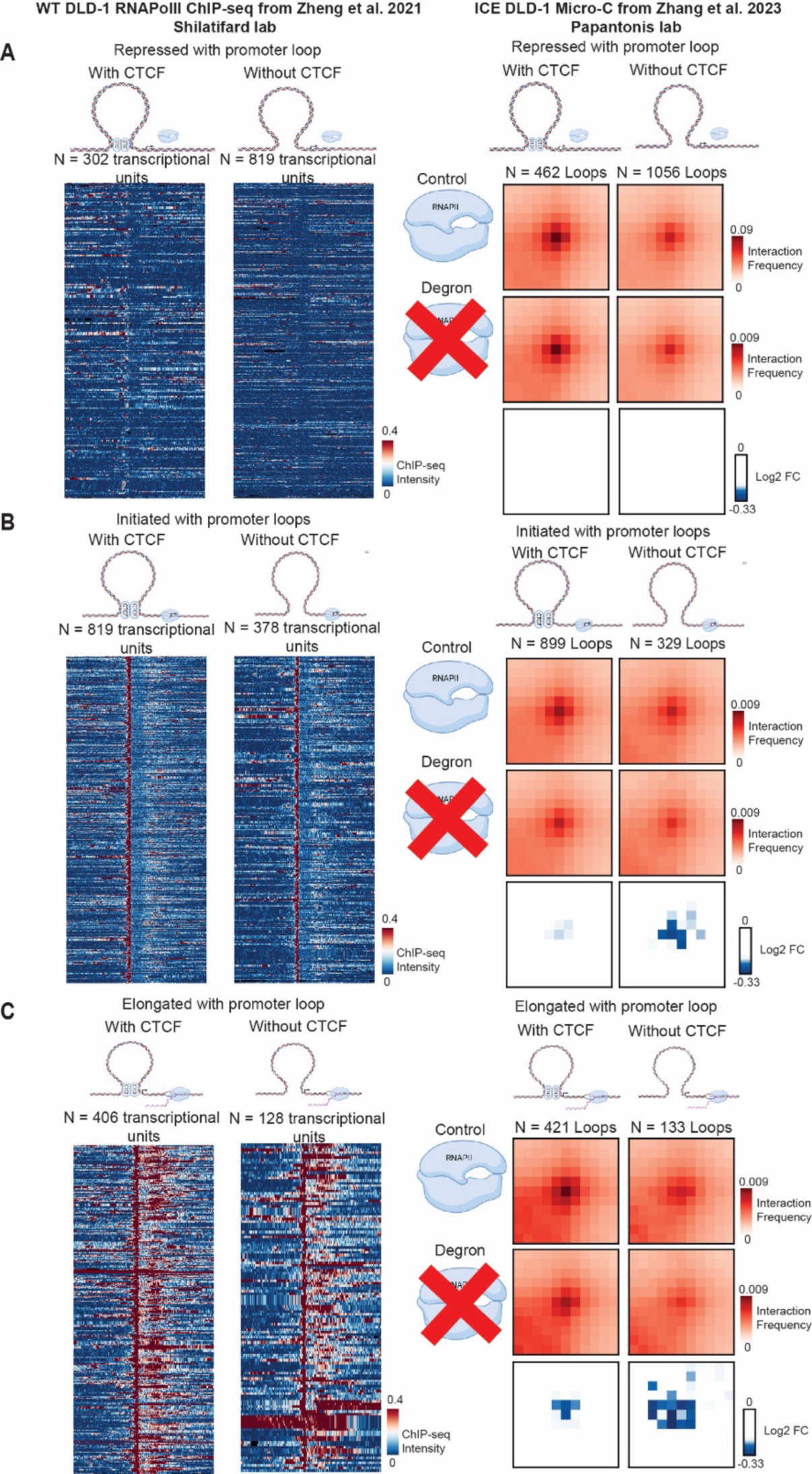
The interaction frequency of loops anchored by elongated genes devoid of CTCF occupancy at their promoter is particularly susceptible to disruption upon RNAPolII degradation. (**A-C**) RNAPolII ChIP-seq heatmaps for transcriptional units categorized as (**A**) repressed, (**B**) initiated, and (**C**) elongated in the wildtype DLD-1 cell line. Transcriptional units are further stratified by CTCF occupancy in a 2 kb window upstream of the TSS. The right column has aggregate peak analysis heatmaps of Micro-C signal at loops called in DLD-1 cells before and after short-term RNAPolII degradation. Some illustrations were created using Biorender.com.

## Discussion

Understanding how chromatin loops interplay with RNAPolII is important toward understanding the principles governing gene expression regulation in development. Here, we use a model of human iPSC differentiation to NPCs and neurons and create genome-wide kb-resolution maps of higher-order chromatin folding, gene expression, and RNAPolII occupancy. We uncover an unexpected strong correlation between cell type-specific E-P loops and genes with RNAPolII occupancy patterns indicative of an initiation-to-elongation transition during differentiation. Elongated genes connected in E-P loops gained de novo during differentiation display robust upregulation of mRNA. By contrast, elongated genes anchoring P-P loops, invariant E-P loops, or unlooped exhibit modest to negligible changes in mRNA levels. Moreover, we observe that genes with RNAPolII patterns indicative of repressed-to-initiated and elongated-to-repressed transitions show changes in mRNA levels whether engaged in loops or not looping. Together our data suggest that robust increases in transcription elongation strongly correlate with de novo E-P loops, whereas transcription repression, initiation, or decommissioning can occur whether the genes engage in loops or not (**Figure 7A**).

**Figure 7.**
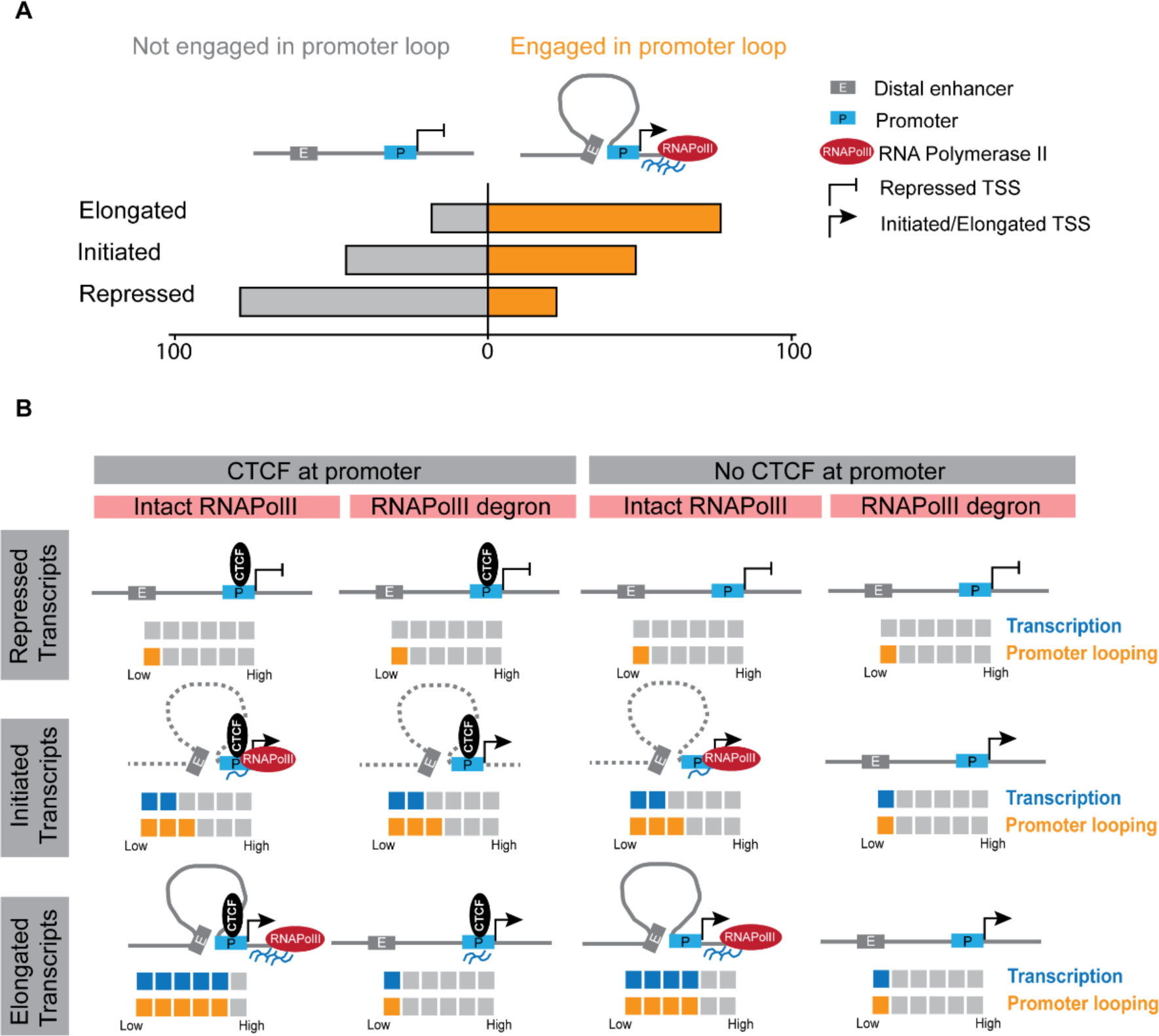
Schematic model of the link among RNAPolII elongation, cell type-specific E-P loops, and the robust upregulation of gene expression during neural cell fate transitions. **(A)** The large majority of genes transitioning from initiated-to-elongated during neural differentiation connect in invariant or cell type-specific looping interactions. Genes that remain repressed during differentiation generally do not loop. Genes transitioning from repressed-to-initiated during differentiation exhibit equal probability of looping or not looping. **(B)** Models representing the sensitivity of loops formed by repressed, initiated, and elongated genes with or without promoter bound CTCF to the short-term degradation of RNAPolII.

The link between RNAPolII and looping has been debated, with conflicting results linked to low-resolution datasets or pharmacological agents with non-specific indirect effects^22,31,32,39–43^. Most recently, two studies employed a degron to achieve short-term RNAPolII degradation and uncovered a previously underappreciated effect on E-P loop maintenance^31,32^. Here, we build on these works by creating genome-wide maps of cell type-invariant and cell type-specific chromatin loops during the transitions from hiPSCs to NPCs and NPCs to post-mitotic neurons. We observe a strong correlation between *de novo* cell type-specific chromatin loops between enhancers and promoters, transcription elongation, and robust upregulation of mRNA levels. By contrast, we find that genes can transition from repressed-to-initiated RNAPII whether they engage in cell type-specific loops, cell type-invariant loops, or do not loop to distal enhancers. These observations are surprising because they support a hypothesized role for E-P loops in the transition from RNAPolII initiation to elongation. Therefore, we re-analyzed published Micro-C data from an established RNAPolII degron system in a dividing mammalian cell line but taking the steps to stratify gene isoforms among those that are repressed, elongated, and initiated. We demonstrate that loops anchored by initiated genes are slightly sensitive to RNAPolII degradation, but only when the promoter is CTCF-independent. CTCF-bound initiated promoters anchor loops that are protected against disruption by RNAPolII knock-down (**Figure 7B**). By contrast, the sensitivity of chromatin loops to RNAPolII degradation is particularly strong when anchored by elongated promoters regardless of CTCF occupancy (**Figure 7B**). Our observations reinforce that RNAPolII causally contributes to the strength of loops anchored by elongated genes.

The models by which transcription start sites (TSSs) communicate with distal CREs to spatiotemporally regulate transcription remain hotly debated when CREs and TSSs are separated by kb to Mb of DNA. One leading model is the formation of stable long-range E-P loops in which promoter activity depends on sustained enhancer proximity via tethering^19,44–47^. A second leading “kiss-and-run” model involves the transient interaction of CREs with their target TSS^48^. Transient E-P contacts could in principle deposit persistent information on the TSSs to facilitate transcription, such as post-translational protein modifications, chromatin modifying enzymes, or Pol II itself. E-P loops would be important for transcription but ensemble Hi-C or DNA FISH measurements of interaction frequency might not correlate with bulk mRNA measurements or even correlate temporally with burst size/frequency^49,50^. A third model, loop-independent long-range communication, involves the diffusion of biomolecules between the CRE and TSS. Finally, a fourth ‘condensate’ model involves the association of promoters and enhancers within foci representing concentrated subnuclear microenvironments of regulatory enzymes, proteins, and RNAs^51^. It has been proposed that Mediator, Brd4, and RNA Pol II can form foci – potentially through liquid-liquid phase-separation mechanisms^52,53^. Here, we take a step toward gathering data to build evidence toward testing these models. We observe that the large majority of elongated genes connect in invariant or cell type-specific looping interactions, whereas repressed genes do not loop and initiated genes exhibit equal probability of looping or not looping (**Figure 7A**). Punctate focal dots represent loops detectable in ensemble Hi-C data because they are typically present in a large proportion of cells and are unlikely to be transient. Therefore, our data are not consistent with transient “kiss-and-run” or loop-independent models for transcription elongation. Rather, our analyses suggest that genes with high levels of RNAPolII elongation and robust upregulation of mRNA levels show genome folding patterns in ensemble Hi-C data more consistent with sustained E-P tethering in loops (model 1). Our data also cannot rule out condensates (model 4). Because condensates range from 200-500 nm in size, they in principle could create local environments of proximal access to similar biomolecules without direct contact^51^. Future causal studies should aim to dissect the causal role for loop extrusion and condensate formation in the loops observed connecting distal enhancers and promoters to transcription elongation.

The models governing long-range E-P communication remain an exciting open area, and further perturbative and single cell imaging and genomics studies will garner further insight toward how each model alone or individually impacts gene regulation at the stages of initiation, pausing, and elongation. Altogether, our results highlight a link between transcriptional elongation and cell type-specific E-P chromatin loops and emphasize the importance of future work dissecting the causal role for looping in gene expression regulation during human neural lineage commitment.

## Author contributions

Conceptualization: KT, HC, ZS, JEPC

Methodology/Visualization: KT, HC, ZS, DP, JEPC

Investigation: KT, HC, ZS, JEPC

Funding: JEPC, DP

Administration: JEPC

Writing & Editing: KT, HC, ZS, JEPC

Reagents: DP

## Data and materials availability

We have provided all data and code to reviewers via private links and will make the links freely available and public upon publication.

