## Supplemental Methods and Figures for "Cell type-specific loops linked to RNA polymerase II elongation in human neural differentiation"

### Supplementary Tables

**Table S1. Long-range looping interactions in human iPSC, NPC, and neuron Hi-C**

**Table S2. Cell type-specific and invariant loop calls for the comparison of human iPSCs and NPCs**

**Table S3. Cell type-specific and invariant loop calls for the comparison of human NPCs to neurons**

**Table S4. Genes classified by RNAPolII occupancy in human iPSCs, NPCs, and neurons**

**Table S5. CTCF ChIP-seq peaks in human iPSCs, NPCs, neurons**

**Table S6. Genes classified by RNAPolII and CTCF occupancy in the DLD-1 cell line**

### Supplementary Figures

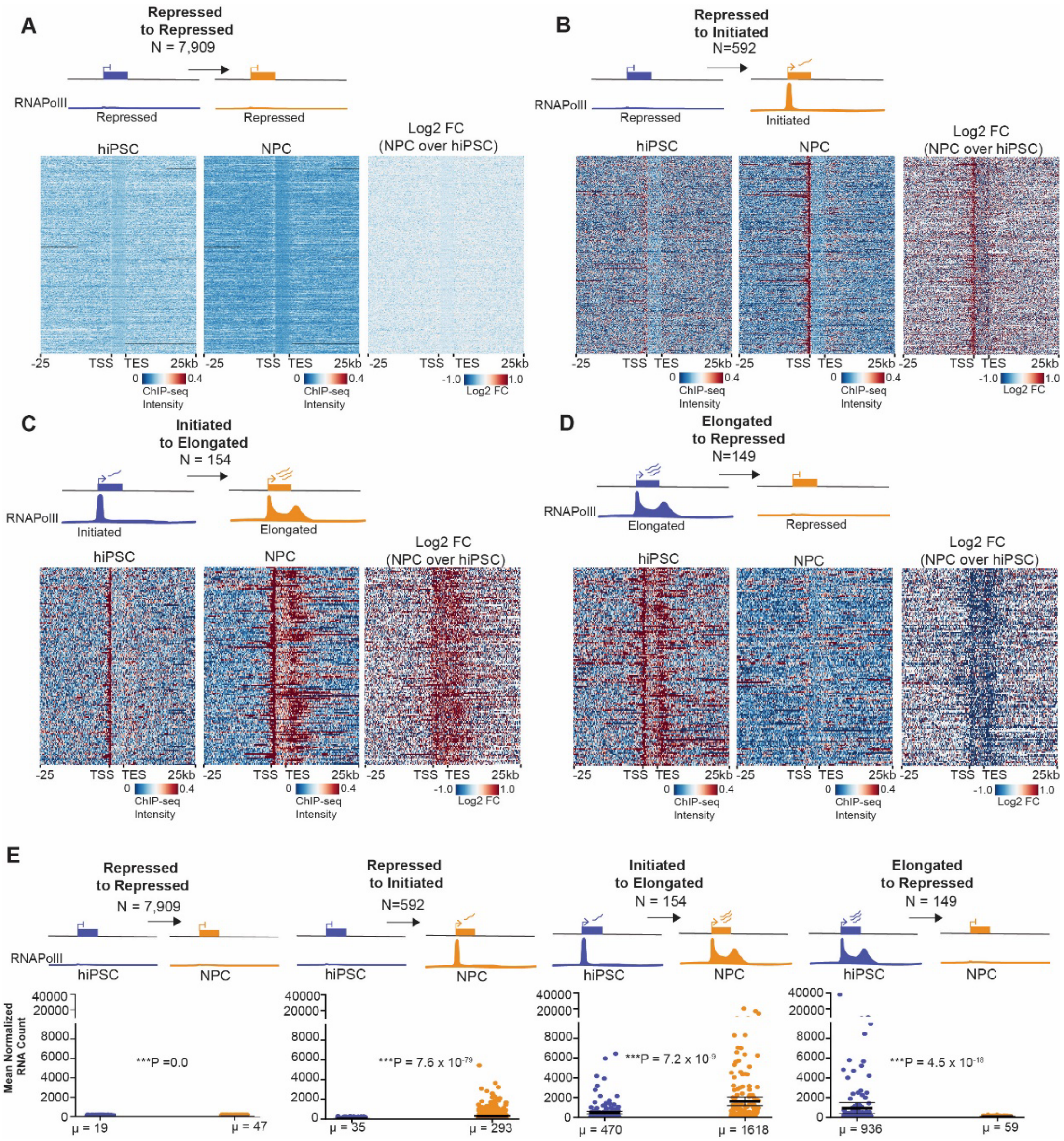

**Supplemental Figure 1. Dynamic RNAPolII occupancy patterns during the cell fate transition from human iPSCs (hIPSCs) to neural progenitor cells (NPCs).**

Related to Figure 3.

(**A-D**) RNAPolII ChIP-seq heatmaps for genes categorized during the transition from iPSC- to NPCs, including **(A)** repressed (hiPSCs) to repressed (NPCs), **(B)** repressed (hiPSCs) to initiated (NPCs), **(C)** initiated (hiPSCs) to elongated (NPCs), and **(D)** elongated (hiPSCs) to repressed (NPCs). Each row represents an individual transcriptional unit. (**E**) mRNA levels from RNA-seq for transcriptional unit classified in A-D. Points represent the mean normalized RNA count from three biological replicates. Horizontal lines represent the mean across all points. P-values, two-tailed Mann Whitney U (MWU) test with significance based on $\alpha = 0.01$.

89,706 transcripts

↓ Filter out transcripts on non-canonical chromosomes

80,377 transcripts

↓ Filter out chrY, bad regions, and small transcripts (< 10kb)

59,750 transcripts

↓ Merge transcripts with the same TSS and TES coordinates

27,030 transcriptional units

↓ Parse transcriptional units into RNAPolII groups

Neuron

| NPC | Repressed | Initiated | Elongated | Unclassified | Total |
|---|---|---|---|---|---|
| Repressed | 9,155 | 612 | 21 | 670 | **9,309** |
| Initiated | 873 | 7,821 | 260 | 1,437 | **9,031** |
| Elongated | 69 | 123 | 1,126 | 358 | **2,962** |
| Unclassified | 415 | 779 | 481 | 2,830 | **5,728** |
| **Total** | **10,512** | **9,335** | **1,888** | **5,295** | **27,030** |

↓ Filter out RNAPolII groups with unclassified categorization

| | Repressed | Initiated | Elongated | Total |
|---|---|---|---|---|
| Repressed | 9,155 | 612 | 21 | **9,788** |
| Initiated | 873 | 7,821 | 260 | **8,954** |
| Elongated | 69 | 123 | 1,126 | **1,318** |
| **Total** | **10,097** | **8,556** | **1,407** | **20,060** |

↓ Parse all classified transcripts into looping classes

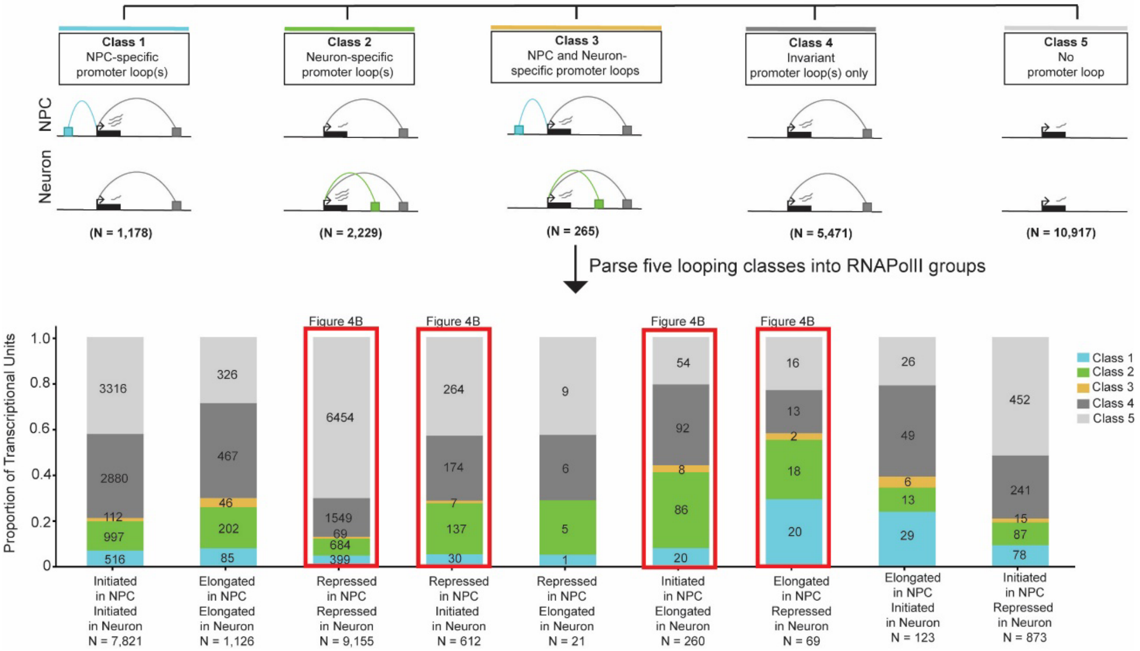

**Supplemental Figure 2. Schematic showing computational steps to stratify genes by RNAPolII occupancy and cell type-specific loops during the cell fate transition of human NPCs to post-mitotic neurons.** Related to Figure 4.

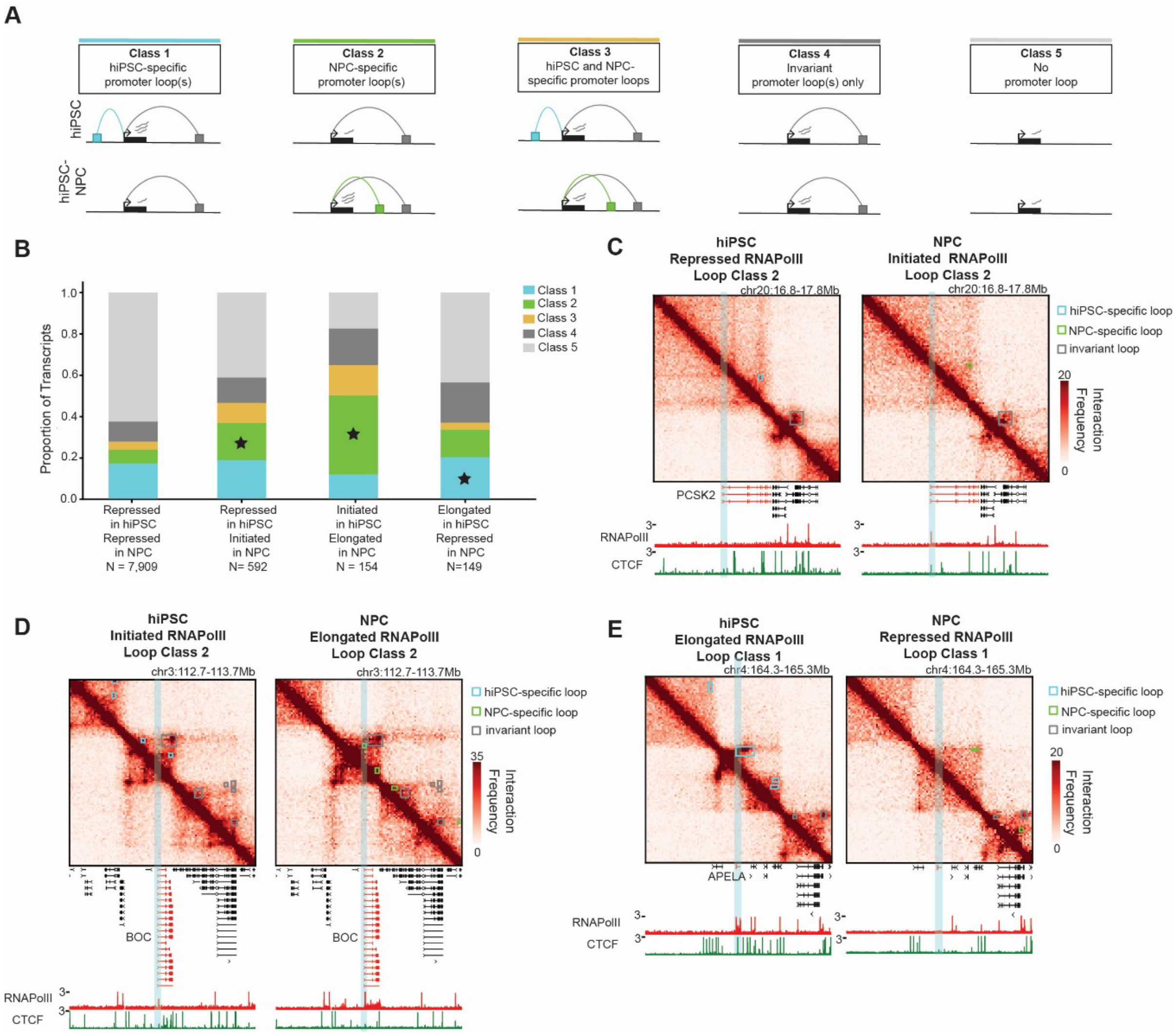

**Supplemental Figure 3. Genes transitioning from initiated to elongated RNAPolII occupancy in NPCs are strongly enriched for NPC-specific loops during differentiation from hiPSCs to NPCs.** Related to Figure 4. (**A**) Schematic depicting promoters of unique transcriptional units classified as anchoring hiPSC-specific loops (Class 1, blue), NPC-specific loops (Class 2, green), mixed loops (Class 3, yellow), cell type-invariant loops (Class 4, dark grey), or not looping (Class 5, light grey). **(B)** Proportion of genes engaged in the five looping categories from panel 4A. Asterisk (*) indicates loop classifications of interest illustrated in panels 4C-E. **(C-E)** Hi-C heatmaps during hiPSC-to-NPC differentiation **(C)** the *PCSK2* gene illustrating RNAPolII occupancy characteristic of repressed to initiated cell fate transition, **(D)** the *BOC* gene illustrating RNAPolII occupancy characteristic of initiated to elongated transition, **(E)** the *APELA* gene illustrating RNAPolII occupancy characteristic of elongated to repressed transition. Tracks below Hi-C heatmaps show RNAPolII and CTCF ChIP-seq data.

89,706 transcripts

↓ Filter out transcripts on non-canonical chromosomes

80,377 transcripts

↓ Filter out chrY, bad regions, and small transcripts (< 10kb)

59,750 transcripts

↓ Merge transcripts with the same TSS and TES coordinates

27,030 transcriptional units

↓ Parse transcriptional units into RNAPolII groups

| hiPSC \ NPC | Repressed | Initiated | Elongated | Unclassified | Total |
|---|---|---|---|---|---|
| Repressed | 7,909 | 592 | 107 | 701 | **9,309** |
| Initiated | 1,470 | 6,776 | 154 | 631 | **9,031** |
| Elongated | 149 | 710 | 1,147 | 956 | **2,962** |
| Unclassified | 930 | 2,313 | 268 | 2,217 | **5,728** |
| **Total** | **10,458** | **10,391** | **1,676** | **4.505** | **27,030** |

↓ Filter out RNAPolII groups with unclassified categorization

| | Repressed | Initiated | Elongated | Total |
|---|---|---|---|---|
| Repressed | 7,909 | 592 | 107 | **8,608** |
| Initiated | 1,470 | 6,776 | 154 | **8,400** |
| Elongated | 149 | 710 | 1,147 | **2,006** |
| **Total** | **9,528** | **8,078** | **1,408** | **19,014** |

↓ Parse all classified transcripts into looping classes

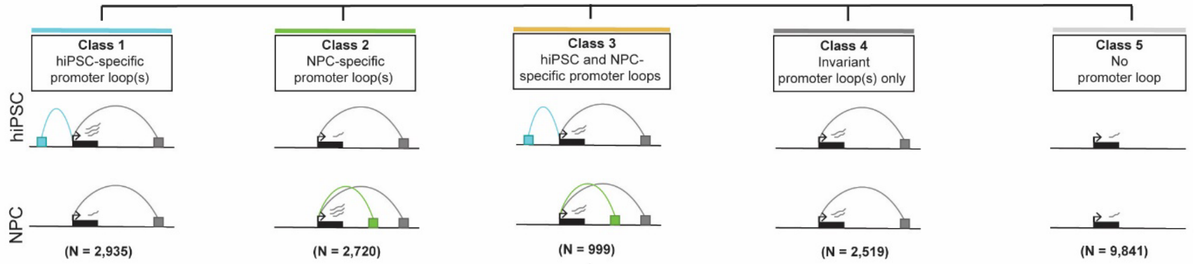

↓ Parse five looping classes into RNAPolII groups

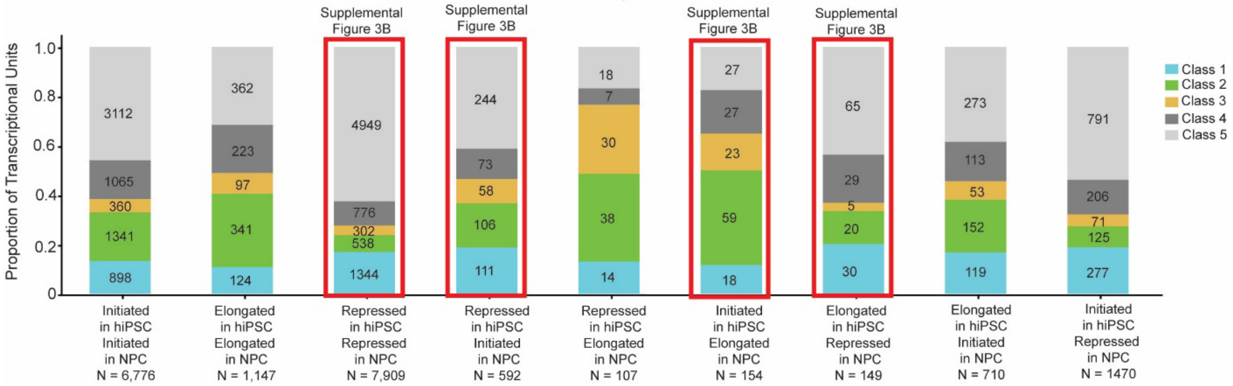

**Supplemental Figure 4. Schematic showing computational steps to stratify genes by RNAPolII occupancy and cell type-specific loops during the cell fate transition of human hiPSCs to NPCs.** Related to Supplemental Figure 3.

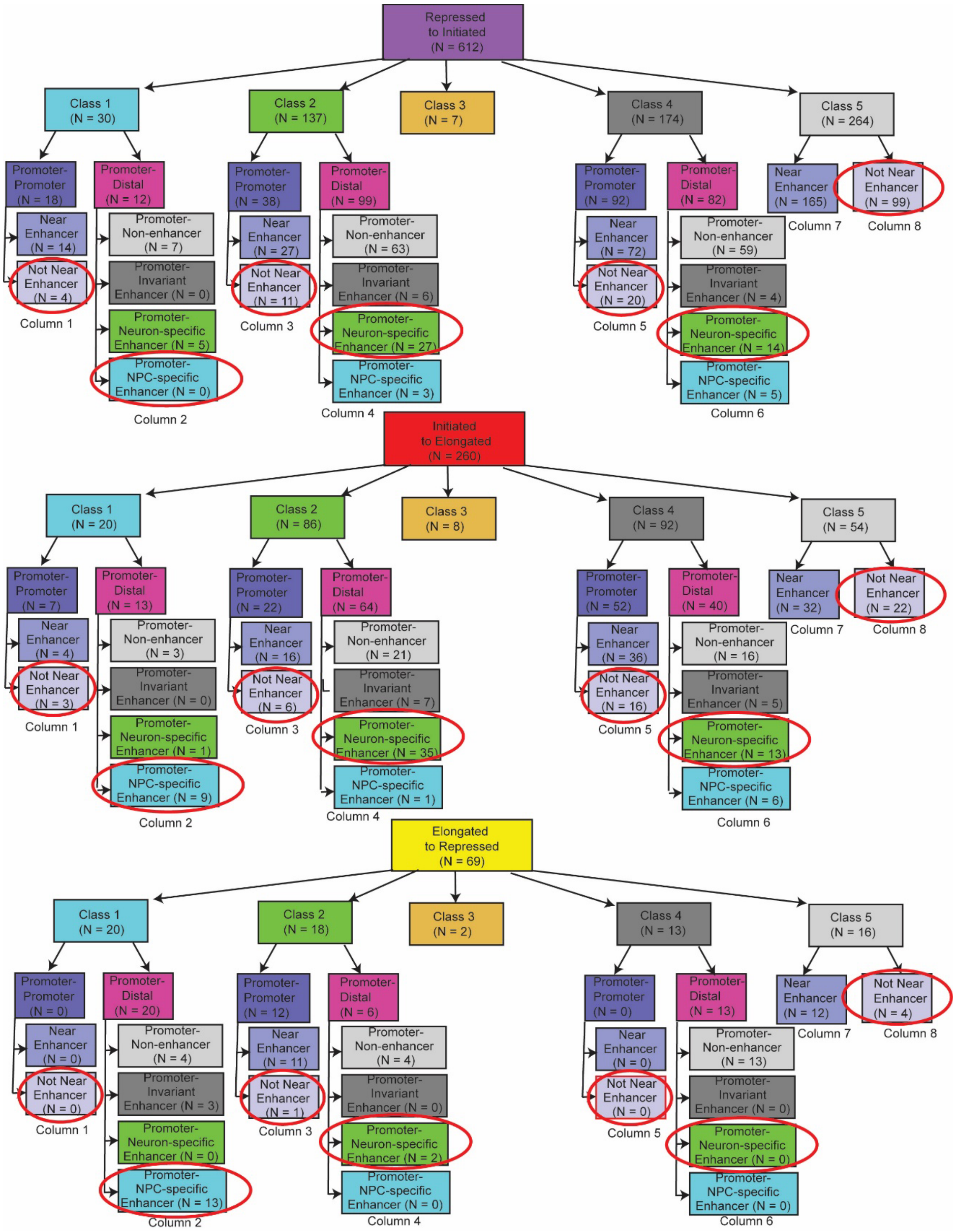

**Supplemental Figure 5**. **Schematic for classifying repressed-to-repressed, repressed-to-initiated, and initiated-to-elongated genes into cell type-specific and invariant enhancer-promoter and promoter-promoter loops during the NPC-to-neuron transition.** Related to Figure 5. Red circles indicate genes analyzed in Figure 5. Rows in Figure 5 include (**A, D**) those groups in red circles transitioning from repressed to initiated RNAPolII occupancy, (**B, E**) those groups in red circles transitioning from initiated to elongated RNAPolII occupancy, (**C, F**) those groups in red circles transitioning from elongated to repressed RNAPolII occupancy during the NPC-to-neuron transition. Rows A-C in Figure 5 include those groups in red circles engaged in (i) promoter to NPC-specific enhancer class 1 NPC-specific loops (column 2), (ii) promoter to neuron-specific enhancer class 2 neuron-specific loops (column 4), (iii) promoter to neuron-specific enhancer class 4 cell type-invariant loops (column 6), (iv) promoters not engaged in loops (column 8). Rows D-F in Figure 5 include those groups in red circles engaged in (i) promoter to promoter class 1 NPC-specific loops (column 1), (ii) promoter to promoter class 2 neuron-specific loops (column 3), (iii) promoter to promoter class 4 cell type-invariant loops (column 5), (iv) promoters not engaged in loops (column 8). To ensure that promoter-promoter loops did not include enhancers, we required that no enhancer was within an 80kb vicinity of the transcription start site (TSS).

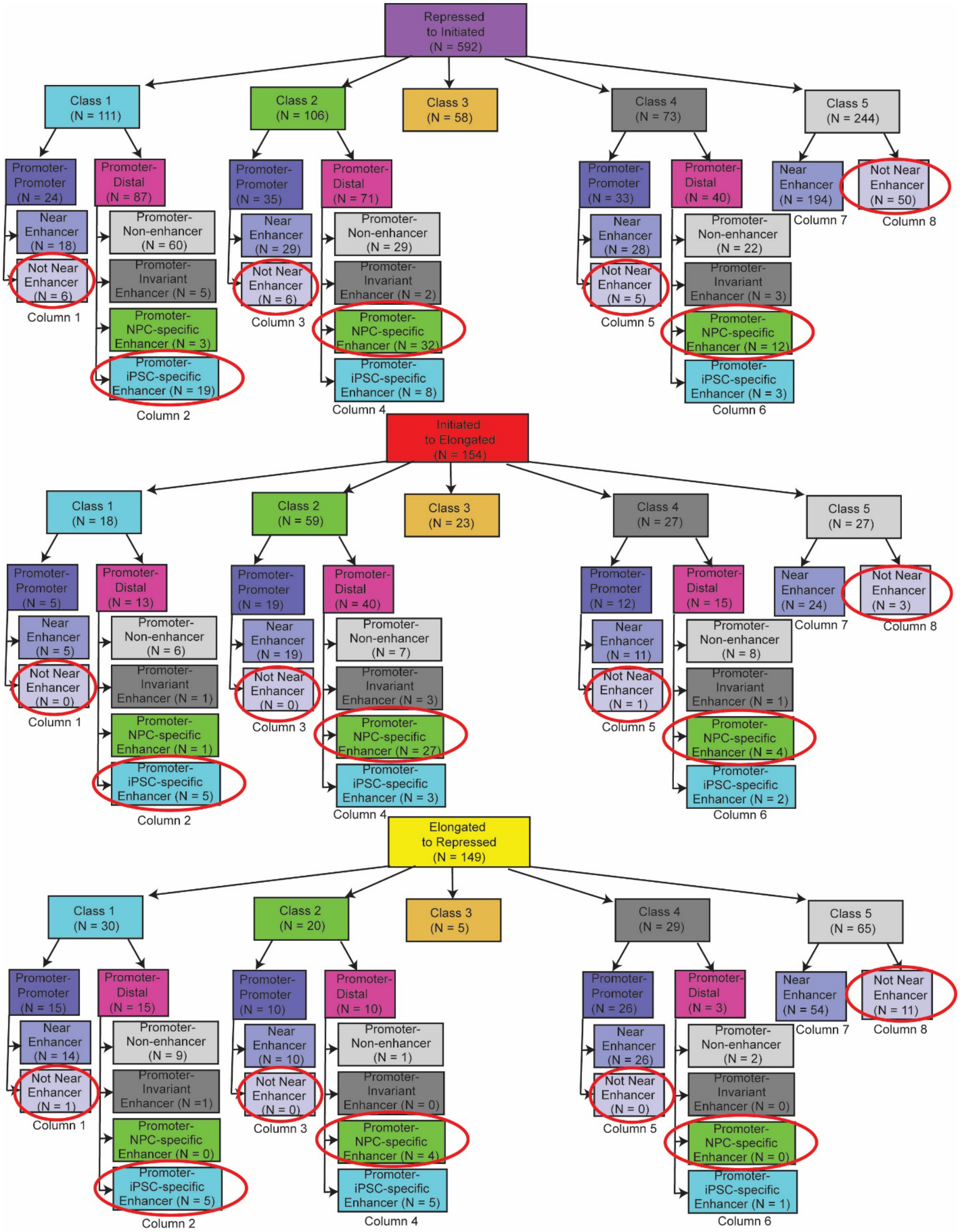

**Supplemental Figure 6. Schematic for classifying repressed-to-repressed, repressed-to-initiated, and initiated-to-elongated genes into cell type-specific and invariant enhancer-promoter and promoter-promoter loops during the hiPSC-to-NPC transition.** Related to

Figure 5 and Figure S7. Red circles indicate genes analyzed in Figure S7. Rows in Figure S7 include (**A, D**) those groups in red circles transitioning from repressed to initiated RNAPolII occupancy, (**B, E**) those groups in red circles transitioning from initiated to elongated RNAPolII occupancy, (**C, F**) those groups in red circles transitioning from elongated to repressed RNAPolII occupancy during the hiPSC to NPC transition. Rows A-C in Figure S7 include those groups in red circles engaged in (i) promoter to hiPSC-specific enhancer class 1 hiPSC-specific loops (column 2), (ii) promoter to NPC-specific enhancer class 2 NPC-specific loops (column 4), (iii) promoter to NPC-specific enhancer class 4 cell type-invariant loops (column 6), (iv) promoters not engaged in loops (column 8). Rows D-F in Figure S7 include those groups in red circles engaged in (i) promoter to promoter class 1 hiPSC-specific loops (column 1), (ii) promoter to promoter class 2 NPC-specific loops (column 3), (iii) promoter to promoter class 4 cell type-invariant loops (column 5), (iv) promoters not engaged in loops (column 8). To ensure that promoter-promoter loops did not include enhancers, we required that no enhancer was within an 80kb vicinity of the transcription start site (TSS).

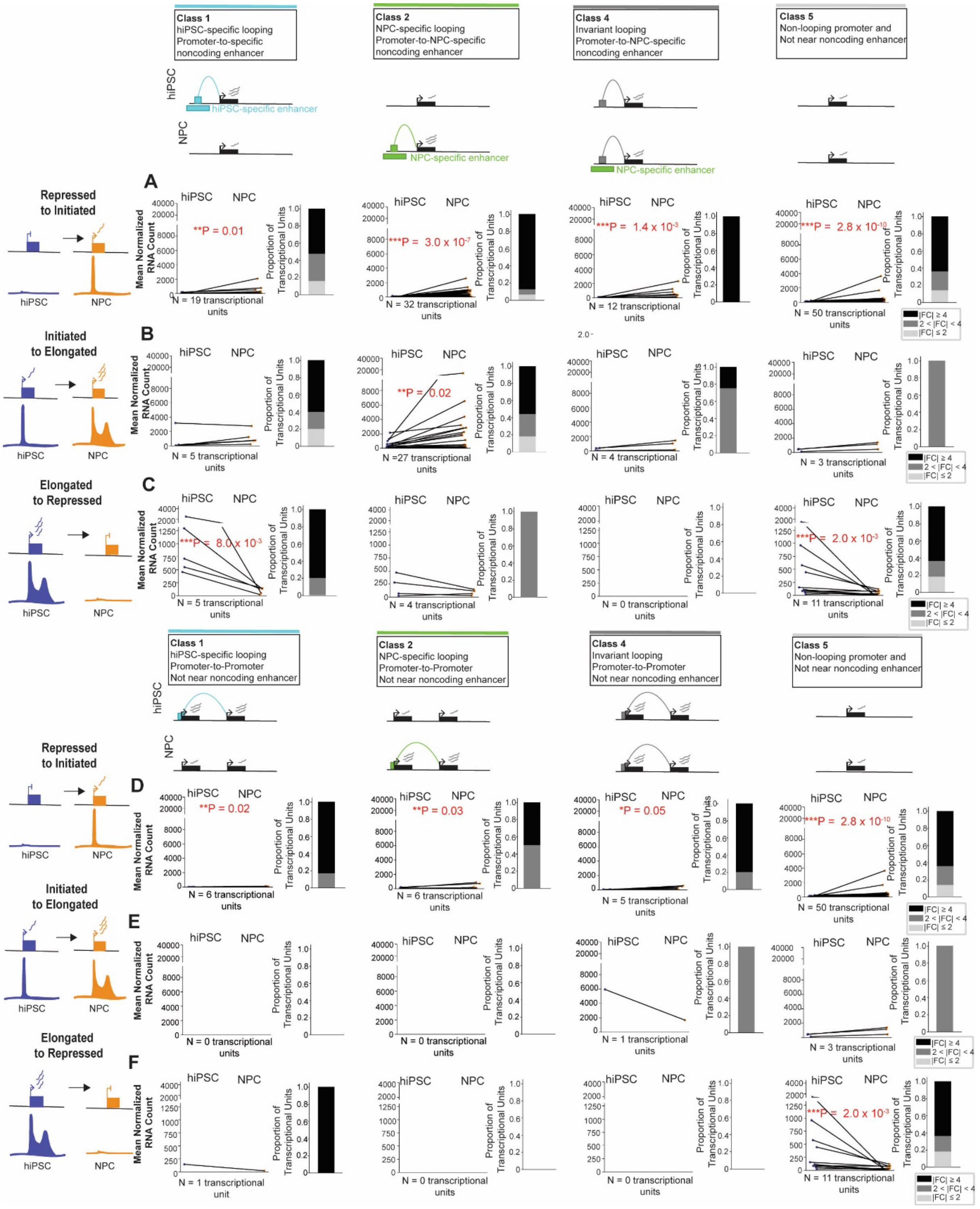

**Supplemental Figure 7. Comparison of mRNA levels for genes engaged in enhancer-promoter loops, promoter-promoter loops, and not looping during hiPSC to NPC differentiation.** Related to Figure S6 and Figure 5.

(**A-F**) mRNA levels calculated from RNA-seq for transcriptional units stratified as (**A, D**) repressed to initiated, (**B, E**) initiated to elongated, and (**C, F**) elongated to repressed during hiPSC to NPC differentiation. (**A-C**) These three gene classes are stratified into those that (column 1) anchor hiPSC-specific enhancer-promoter loops decommissioned in differentiation (Class 1, blue), (column 2) anchor NPC-specific enhancer-promoter loops gained de novo in differentiation (Class 2, green), (column 3) anchor cell type-invariant loops connecting promoters to NPC-specific promoters (Class 4, dark grey), and (column 4) do not loop (Class 5, light grey). (**D-F**) These three gene classes are stratified into those that (column 1) anchor hiPSC-specific promoter-promoter loops decommissioned in differentiation (Class 1, blue), (column 2) anchor NPC-specific promoter-promoter loops gained de novo in differentiation (Class 2, green), (column 3) anchor cell type-invariant promoter-promoter loops (Class 4, dark grey), and (column 4) do not loop and lack enhancers within a 80kb vicinity of the TSS (Class 5, light grey). Each point is the mean normalized RNA count across three replicates. P-values are computed using a two-tailed Mann Whitney U test. Asterisk (*) reach significance under $\alpha = 0.1$ and triple asterisks (***) reach significance under $\alpha = 0.01$. Stacked bar plots represent the proportion of genes that exhibit absolute fold-change expression less than 2, greater than 2 and less than 4, and greater than 4 between hiPSC and NPC cellular states. Groups with less than 10 transcriptional units in panels A-C are greyed out due to the low number of genes.

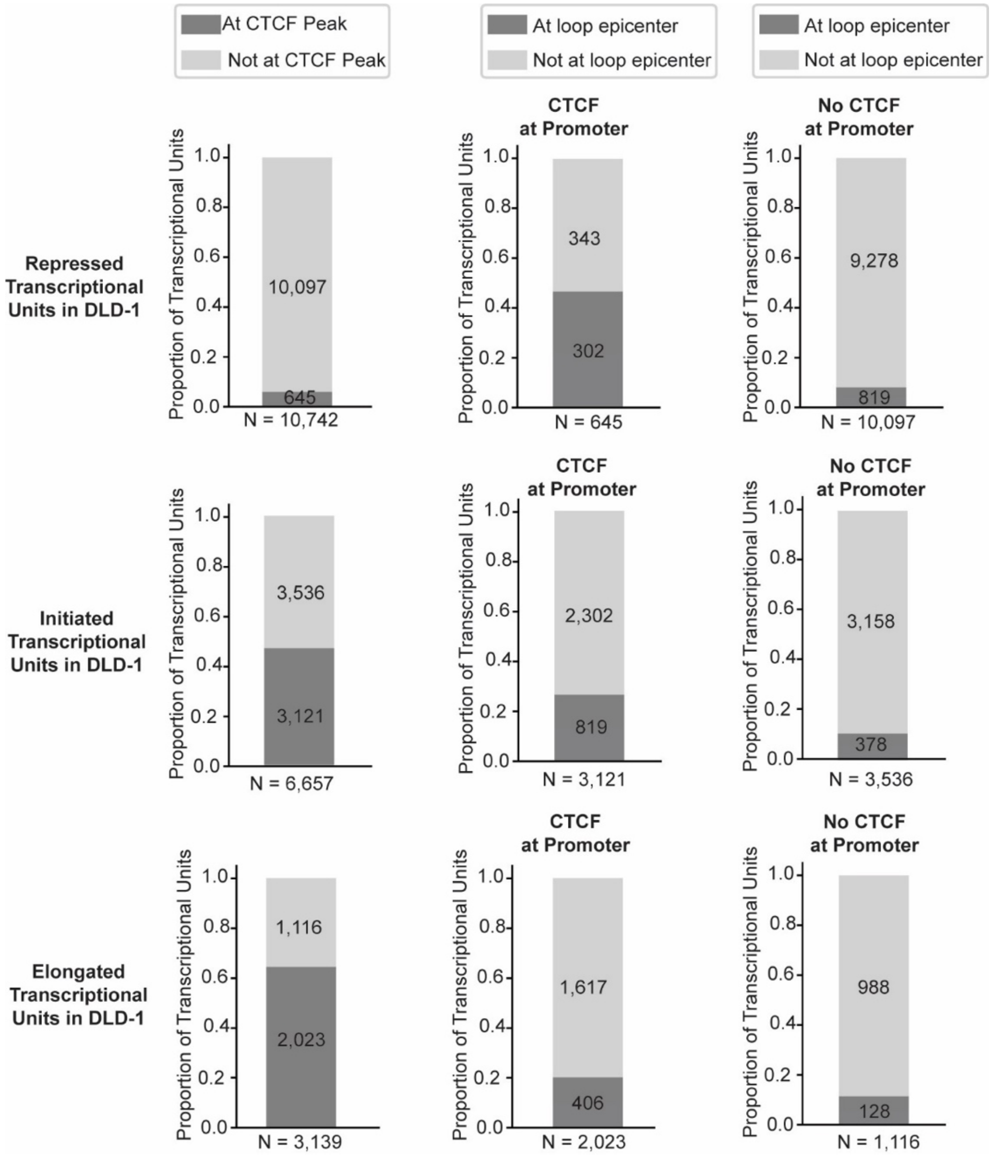

**Supplemental Figure 8. Genes in DLD-1 cells classified by RNAPolII occupancy patterns characteristic of repression, initiation, and elongation and with or without CTCF occupancy at their promoters.** Related to Figure 6.

Column 1, the proportion of repressed, initiated, and elongated transcriptional units genome-wide with or without CTCF binding at their promoters in DLD-1 cells. Column 2, the proportion of repressed, initiated, and elongated transcriptional units genome-wide with CTCF binding at their promoters that are also anchoring the base of loops as detected by Micro-C in DLD-1 cells. Column 3, the proportion of repressed, initiated, and elongated transcriptional units genome-wide without CTCF binding at their promoters that are also anchoring the base of loops as detected by Micro-C in DLD-1 cells.

### STAR METHODS

#### KEY RESOURCE TABLE

| REAGENT or RESOURCE | SOURCE | IDENTIFIER |
|---|---|---|
| Antibodies | | |
| OCT4 | Santa Cruz Biotechnology | Cat# SC9081 , RRID:AB_2167703 |
| SSEA4 | R&D Systems | Cat# MAB1435, RRID:AB_357704 |
| Nestin | R&D Systems | Cat# MAB1259, RRID:AB_2251304 |
| FoxG1 | Abcam | Cat# ab18259, RRID:AB_732415 |
| CTIP2 | Abcam | Cat# ab18465, RRID:AB_2064130 |
| SATB2 | Abcam | Cat# ab51502, RRID:AB_882455 |
| Anti-Mouse IgG H&L Alexa Fluor® 488 | Abcam | Cat# ab150113, RRID:AB_2576208 |
| Anti-Rat IgG H&L Alexa Fluor® 647 | Abcam | Cat# ab150159, RRID:AB_2566823 |
| Goat Anti-Rabbit IgG H&L Alexa Fluor® 594 | Abcam | Cat# ab150080, RRID:AB_2650602 |
| IgG from rabbit serum | Sigma-Aldrich | Cat# I8140, RRID:AB_1163661 |
| CTCF | Millipore | Cat# 07-729, RRID:AB_441965 |
| H3K27ac | Millipore | Cat# 07-449, RRID:AB_310624 |
| YY1 | Abcam | Cat# ab109237, RRID:AB_10890662 |
| RNA pol II antibody (mAb) (Clone 4H8) | Active Motif | Cat# 39097, RRID:AB_2732926 |
| Chemical, peptides, and recombinant proteins | | |
| StemFlex culture media | Thermo Fisher Scientific | Cat # A3349401 |
| Geltrex | Thermo Fisher Scientific | Cat # A1569601 |
| Essential 6 media | Thermo Fisher Scientific | Cat # A1516401 |
| Penicillin-streptomycin | Gibco | Cat# 15140122 |
| Versene Solution | Gibco | Cat# 15040066 |
| Accutase | Gibco | Cat# A1110501 |
| Y-27632 (Dihydrochloride) | Stem Cell Technologies | Cat# 72304 |

| | | |
|---|---|---|
| Neurobasal media | Thermo Fisher Scientific | Cat# 21203-049 |
| DMEM/F12 GlutaMax | Thermo Fisher Scientific | Cat# 10565-018 |
| B27 supplement | Thermo Fisher Scientific | Cat# 17504-044 |
| GlutaMax supplement | Thermo Fisher Scientific | Cat# 35050 |
| Non-Essential Amino Acid | Thermo Fisher Scientific | Cat# 11140-050 |
| N-2 supplement | Thermo Fisher Scientific | Cat# 17502048 |
| Insulin | Sigma | Cat# I1882 |
| 2-mercapto-ethanol | Thermo Fisher Scientific | Cat# 21985-023 |
| SB431542 | Selleckchem | Cat# S1067 |
| LDN-193189 | Selleckchem | Cat# S2618 |
| poly-L-ornithine | Sigma-Aldrich | Cat# P4957 |
| Laminin | Thermo Fisher Scientific | Cat# 23017015 |
| FGF-2 | Stem Cell Technologies | Cat# 78003 |
| Uridine | Sigma | Cat# U3750-1G |
| STEMdiff Neural Rosette Selection Reagent | Stem Cell Technologies | Cat# 5832 |
| DAPT | Sigma-Aldrich | Cat# D5942 |
| PBS | Corning | Cat# 21-040-CV |
| 100% Ethanol | Decon Labs | Cat# 2716 |
| EDTA, pH 8.0 | Invitrogen | Cat# 15575020 |
| Synth-a-Freeze | Gibco | Cat# A1254201 |
| Formaldehyde solution | Sigma-Aldrich | Cat# F8775 |
| Formaldehyde solution | Pierce | Cat# 28908 |
| HEPES-KOH, pH 7.5 | Boston BioProducts | Cat# BBH-75-K |
| Igepal CA-630 | Sigma-Aldrich | Cat# I8896 |
| 5M NaCl | Invitrogen | Cat# AM9760G |
| Protease Inhibitor | Sigma | Cat# P8340 |
| SDS solution, 10% | Fisher Scientific | Cat# 15553027 |
| TWEEN 20 | Sigma | P9416-50ML |
| Sodium bicarbonate | Sigma-Aldrich | Cat# S5761 |
| Sodium deoxycholate | Sigma-Aldrich | Cat# D6750 |
| TE buffer, pH 8.0 | Invitrogen | Cat# AM9858 |
| Lithium chloride, ultra dry | Alfa Aesar | Cat# 1368406 |
| Magnesium chloride solution | Sigma | Cat# M1028-100ML |
| Tris-HCl, pH 8.0 | Invitrogen | Cat# 15568025 |
| Triton X-100 solution | Sigma-Aldrich | Cat# 93443 |
| TE buffer, pH 8.0 | Invitrogen | Cat# AM9858 |
| Nuclease-free water | Sigma-Aldrich | Cat# W4502 |
| VECTASHIELD Antifade Mounting Medium | Vector Laboratories | Cat# H-1200 |
| Protein A Agarose beads | Invitrogen | Cat# 15918014 |
| Protein G Agarose beads | Invitrogen | Cat# 15920010 |
| Agencourt AMPure XP beads | Beckman Coulter | Cat# A63881 |
| RNaseA | Roche | Cat# 10109169001 |
| Proteinase K | NEB | Cat# P8107S |

| | | |
|---|---|---|
| PMSF solution | Sigma | Cat# 93482-50ML-F |
| Nuclease-free water | Sigma-Aldrich | Cat# W4502 |
| Ultrapure Phenol/Chloroform/Isoamyl Alcohol | Fisher Scientific | Cat# BP1752I100 |
| Critical commercial assays | | |
| Human Karyotype Panel | NanoString | Cat# XT-CSO-KAR15-012 |
| Mycoplasma detection kit | ATCC | Cat# 30-1012K |
| Direct-zol total RNA isolation kit | Zymo | Cat# R2061 |
| Qubit RNA HS assay | Invitrogen | Cat# Q32852 |
| RNA 6000 Pico reagent kit | Agilent Technologies | Cat# 5067-1511 |
| TruSeq® Stranded Total RNA Library Prep Human/Mouse/Rat (48 Samples) (gold) | Illumina | Cat# 20020598 |
| TruSeq RNA Single Indexes Set A and B | Illumina | Cat# 20020492 |
| Qubit dsDNA HS assay kit | Invitrogen | Cat# Q32851 |
| Agilent DNA 1000 reagent kit | Agilent Technologies | Cat# 5067–1504 |
| Arima-HiC kit | Arima Genomics | Cat# A510008 |
| Kapa Library Quantification Kit | KAPA Biosystems | Cat# KK4835 |
| NEBNext Ultra II DNA Library Prep Kit for Illumina | New England Biolabs | Cat# E7645S |
| NextSeq® (150-cycle) v2.5 High Output kit | Illumina, Inc | Cat# 20024907 |
| NextSeq® 500 High Output v2 Kit (75 cycles) kit | Illumina, Inc | Cat# 20024906 |
| Deposited data | | |
| RNAPolII ChIP-seq in hiPSC, hiPSC-NPC, and NPC-Neuron | This study | GEO: GSE220103 |
| CTCF ChIP-seq in hiPSC, hiPSC-NPC, and NPC-Neuron | This study | GEO: GSE220103 |
| H3K27ac ChIP-seq in hiPSC, hiPSC-NPC, and NPC-Neuron | This study | GEO: GSE220103 |
| Hi-C in hiPSC, hiPSC-NPC, and NPC-Neuron | This study | GEO: GSE220103 |
| RNA-seq in hiPSC, hiPSC-NPC, and NPC-Neuron | This study | GEO: GSE220103 |
| Experimental models: cell lines | | |
| 7889SA3.5 | Paquet and colleagues[1] | |
| Software and algorithms | | |
| ImageJ | NIH | https://imagej.nih.gov/ij/ |

| | | |
|---|---|---|
| Adobe Photoshop (v 23.2.2) | Adobe | https://www.adobe.com/ |
| Leica LAS X | Leica | https://www.leica-microsystems.com/products/microscope-software/p/leica-las-x-ls/downloads/ |
| deeptools (v 3.5.1) | [2] | https://deeptools.readthedocs.io/en/develop/ |
| MACS2 (v 2.1.1.20160309) | [3] | https://pypi.org/project/MACS2/ |
| Bowtie (v 0.12.7) | [4] | http://bowtie-bio.sourceforge.net/index.shtml |
| Bedtools (v 2.29.1) | [5] | https://github.com/arq5x/bedtools2 |
| HiC-Pro (v 2.7.7) | [6] | https://github.com/nservant/HiC-Pro |
| DESEQ2 (v 1.22.1) | [7] | doi: 10.1186/s13059-014-0550-8 |
| Kallisto (v 0.46.1) | [8] | https://pachterlab.github.io/kallisto/about |
| tximport | [9] | https://bioconductor.org/packages/release/bioc/html/tximport.html |
| WebGestalt (v 0.4.4) | [10] | https://github.com/bzhanglab/WebGestaltR |
| GEM-mappability (Binary pre-release 2) | [11] | https://sourceforge.net/projects/gemlibrary/files/gem-library/ |
| 3DeFDR-HiC (v 0.2.1) | [12] | https://bitbucket.org/creminslab/hic3defdr |
| pyBigWig (v 0.3.18) | [2] | https://github.com/deeptools/pyBigWig |
| opencv-python (v 4.5.4.58) | [13] | https://docs.opencv.org/4.x/d6/d00/tutorial_py_root.html |
| Mustache (v 1.3.1) | [14] | https://github.com/ay-lab/mustache |

### METHODS DETAILS

#### *Maintenance of human induced Pluripotent Stem Cell (hiPSC) culture*

A subclone of the previously described 7889SA human induced pluripotent stem cell (hiPSC) line was used for each experiment. The line was established from the AG07889 male fibroblast (Coriell Institute for Medical Research, Camden, NJ) and previously characterized and described[1]. Upon arrival, the subclone (7889SA3.5) was expanded and a master stock of cells were frozen from the early passage. The same master stocks were thawed, cultured, and subsequently used for each experiment.

hiPSC cells were cultured on 10 cm cell culture plastic dishes (Corning, #430167), pre-coated with 6 mL Geltrex hESC-Qualified Reduced Growth Factor Basement Membrane Matrix (Thermo Fisher Scientific, #A1569601) for 1 hour at 37 °C. hiPSCs were plated and maintained in StemFlex stem cell culture media (Thermo Fisher Scientific, #A3349401) supplemented with 1 % (v/v) penicillin-streptomycin (Thermo Fisher Scientific, #15140122). Cell cultures were kept in humidified incubator at 37 °C and 5 % $CO_2$. To passage, cells were detached from dishes by washing with 1xPBS and then incubating with 4 ml of Versene Solution (Thermo Fisher Scientific, 15040066) at 37 °C for 3-5 minutes. Following inactivation by the addition of 10 ml of fresh cell culture media, cells were split in a 1:3 to 1:8 dilution and seeded onto Geltrex-coated plates, prepared freshly on the same day.

To verify the cellular state of the hiPSC clones, an early passage stock vial was characterized for (1) normal karyotype, (2) characteristic stem cell morphology, (3) expression of genes indicating pluripotent state, and (4) absence of mycoplasma. A human Karyotype Panel (NanoString, XT-CSO-KAR15-012) was used to assess karyotype from genomic DNA isolated from the master stock. DNA was extracted from the cells using phenol:chloroform and ethanol precipitation and submitted for NanoString service provided by the Genomics Facility at The Wistar Institute, Philadelphia, PA. Immunofluorescence staining was used to characterize the presence of pluripotency markers, we used for OCT4 (Santa Cruz, SC9081, 1:100 dilution) and SSEA4 (R&D, MAB1435, 10 μg/mL) as described in detail under "Immunocytochemistry and microscopy". Absence of mycoplasma contamination was confirmed by collecting $10^5$ hiPSCs into the existing culture media and processing the sample using a mycoplasma detection kit (ATCC, 30-1012K), according to the manufacturer's protocol. Following the initial characterization of the master stock, cultures were continuously monitored for expected pluripotent cell morphology by daily visual assessment under the microscope and routine staining for OCT4 and SSEA4.

***hiPSC differentiation to neural progenitors (NPCs) and post-mitotic neurons***

NPCs and post-mitotic neurons were differentiated from hiPSCs using our recently described cortical neuron differentiation protocol, which reproducibly produces high yields of pure NPCs and neurons, with a few additional optimizations[15]. Briefly, 24 hours prior to starting differentiation, StemFlex media was replaced with Essential 6 (E6) media (Thermo Fisher Scientific, A1516401) on hiPSCs growing on 10 cm cell culture dishes. Induction of neural precursor cell differentiation was initiated by detaching hiPSCs with 4ml Accutase (Thermo Fisher Scientific, #A1110501) followed by Accutase inactivation and single cell suspension by triturating cells with 16 ml of E6 media. Cells were centrifuged, resuspended in E6 media supplemented with ROCK inhibitor (Stem Cell Technologies, #72304) and seeded into Geltrex-coated 12-well tissue culture plates (Falcon, #353043), Medium was replaced to 2 mL neural induction (NI) medium supplemented with LDN-193189 (Selleckchem, #S2618, 1:10000) and SB431542 (Selleckchem, #S1067, 1:1000). The day of change to NI media was counted as day *in vitro* 0 (DIV0). Cells were maintained in NI media for 8 days (DIV8), changing media every day. To differentiate hiPSC derived neural precursor cells (hiPSC-NPCs), DIV8 cells were enzymatically dissociated using Accutase and pelleted cells resuspended at 30 million cells per ml in NI medium, supplemented with ROCK inhibitor. Cells were plated as 250-µl spots onto dry 6-well plates, pre-coated with poly-L-ornithine (Sigma-Aldrich, #P4957) and laminin (Thermo Fisher Scientific, #23017015, 20x dilution), and 2 ml NI medium supplemented with ROCK inhibitor were slowly added to avoid detachment of the cells. Media was daily changed until DIV10. On DIV10 NI media was replaced with neural maintenance (NM) media. 20 ng ml FGF-2 (Stem Cell Technologies Inc., #78003) were added for 2 days as soon as neural rosettes were apparent in the cultures. When cells with neuronal morphology started to emerge from rosettes, rosettes were manually isolated under the microscope under sterile conditions and cultivated in NM media until DIV35, when NPC samples were harvested.

To further differentiate cells into terminally differentiated, post-mitotic neurons (NPC-Neurons), NPC rosettes were triturated into single cells using Accutase, followed by inactivation with fresh media (Neurobasal medium supplemented with B-27 serum-free supplement (Thermo

Fisher Scientific, #17504044), 2 mM GlutaMax (Thermo Fisher Scientific, #35050061), 1% (v/v) Penicillin-Streptomycin (Thermo Fisher Scientific, #15140-122)). Cells were seeded at a ~200,000–500,000 hiPSC-NPCs per well density onto a 24-well poly-L-ornithine/laminin-coated plate. In the first 7 days after plating, 10 µM DAPT (Sigma-Aldrich, #D5942) and 5-FU was added to the media to eliminate proliferating cells and enhancing terminal differentiation. Neurons were harvested at DIV65. Homogenous morphology was confirmed, and presence of neuronal markers were assessed as described in detail under "Immunocytochemistry and microscopy".

***Immunocytochemistry and microscopy***

Stem cell markers (OCT4 and SSEA4), neural precursor cell markers (NESTIN, FOXG1), and mature neuronal markers (CTIP2, SATB2) were assessed by immunofluorescence staining at Day 0, DIV35 (hiPSC-NPC), and DIV65 (NPC-Neuron). Briefly, cells were fixed in 4% paraformaldehyde (Pierce, cat# 28908), permeabilized in PBS/0.1% Triton X-100, and stained with primary antibodies anti-OCT4 (Santa Cruz, #SC9081, 1:100 dilution), anti-SSEA4 (R&D, #MAB1435, 10 µg/mL), anti-FoxG1 (Abcam, #ab18259), anti-Nestin (R&D System, #MAB1259), anti-SATB2 (Abcam, #ab51502) and anti-CTIP2 (Abcam, #ab18465). Secondary antibodies Goat Anti-Mouse IgG H&L Alexa Fluor® 488 (Abcam, #ab150113), Goat Anti-Rat IgG H&L Alexa Fluor® 647 (Abcam, #ab150159), and Goat Anti-Rabbit IgG H&L Alexa Fluor® 594 (Abcam, #ab150080) were used for visualization. Cells were imaged on a Leica DMi8 microscope using software Leica LAS X and images were processed in ImageJ and Adobe Photoshop (Adobe). Image processing was limited to altering brightness and contrast levels equivalently across all images to obtain higher quality phase-contrast images. Parameters were selected after ensuring that secondary antibody only controls did not show signal.

***Total RNA-seq***

Total RNA was extracted from 3 biological replicates of hiPSCs, NPCs and neurons using the Direct-zol total RNA isolation kit (Zymo, #R2061) according to the manufacturer's protocol. Integrity of the RNA samples (RIN) were assessed using Agilent RNA 6000 Pico reagent kit on the Bioanalyzer 2100 (Agilent Technologies, Santa Clara, CA, USA). 500 ng of isolated total RNA with a RIN value above 8 was used for ribosomal RNA depleted strand-specific RNA library preparation, using the TruSeq Stranded Total RNA LT sample preparation kit with Ribo-Zero Gold

(Illumina, #RS-122-2301). TruSeq RNA Single Indexes Set A and B were used and ligated onto cDNA (Illumina, #20020492) to enable multiplex sequencing. Library clean-up and size selection of ~300 bp fragments was performed by Agencourt AMPure XP beads (Beckman Coulter, #A63881) prior to PCR amplification and index incorporation for 15 cycles. Library quality and quantities were assessed using the Qubit high sensitivity DNA assay kit (Thermo Fisher Scientific, Q32851) and the Agilent DNA 1000 reagent kit (Agilent Technologies, 5067–1504) on the Agilent Bioanalyzer 2100 (Agilent Technologies, Santa Clara, CA, USA). Sequencing of multiplexed, pooled samples were performed on Illumina NextSeq500 using a 150-cycle v2.5 High Output kit (Illumina, Cat# 20024907), generating 2 x 75bp, paired-end reads.

***Chromatin fixation for ChIP-seq and Hi-C***

Cells were fixed for downstream ChIP-seq and Hi-C assays as previously described[12,16-22]. Briefly, media was discarded, cells were washed with 6 mL of 1x PBS and 10 mL of freshly prepared fixation media (1 % (v/v) formaldehyde in DMEM/F-12 (Thermo Fisher Scientific, #11320033)) was added to the cells growing on 10 cm cell culture dishes and incubated for 10 min at room temperature. Fixation media was diluted from 11% formaldehyde solution (50 mM HEPES-KOH (pH 7.5), 100 mM NaCl, 1 mM EDTA, 0.5 mM EGTA, 37% formaldehyde (Sigma, Cat# F8775). The fixation process was terminated by adding 125 mM glycine for 5 min at room temperature, followed by 15 min incubation at 4°C. Cross-linked cells were washed with pre-chilled PBS twice before flash freezing in liquid nitrogen. Pellets of cells were stored at −70°C.

***ChIP-seq***

ChIP-seq libraries were prepared as previously described[12,16-22], with minor modifications. Briefly, crosslinked cell pellets (8-10 million cells) were lysed in pre-chilled cell lysis buffer (10 mM Tris pH 8.0, 10 mM NaCl, 0.2 % (v/v) NP-40, 1 mM PMSF, 0.2 % (v/v) Protease inhibitor cocktail (Sigma, #P8340)) on ice for 10 min. Cells were homogenized using a dounce homogenizer and 25-30 strokes with pestle A. To pellet and solubilize nuclei, lysates were centrifuged at 2,500 x g for 5 min at 4°C and pellets were resuspended in 500 µl nuclear lysis buffer (50 mM Tris pH 8.0, 10 mM EDTA, 1 % (w/v) SDS, 1 mM PMSF, 0.2 % (v/v) Protease inhibitor cocktail) on ice for 20 min before mixing with 300 µl IP dilution buffer (20 mM Tris pH 8.0, 2 mM EDTA, 150 mM NaCl, 1 % (v/v) Triton X-100, 0.01 % (w/v) SDS, 1 mM PMSF, 0.2 % Protease inhibitor cocktail).

Chromatin was sheared to ~200-600 bp DNA fragment size using Qsonica Q800R3 (Qsonica Sonicators, CT) with parameters of 100% amplitude and 30 sec on / 30 sec off pulses. Fragment size was confirmed on 1% agarose gel from purified input samples. After sonication lysates were centrifuged at 16,000 x g and incubated for 2 hours with pre-clearing solution consisting 50 µg of IgG (Sigma, #I8140), 175 µl of Protein A Agarose (Thermo Fisher Scientific, #15918014), and 175 µl of Protein G Agarose (Thermo Fisher Scientific, #15920010) in 3.7 ml of pre-chilled IP dilution buffer and 0.5 ml of nuclear lysis buffer by rotating at 10 rpm at 4°C. Chromatin immunoprecipitation was performed from supernatant using antibody-bound beads by rotating the samples at 10 rpm overnight (~12 hours) at 4°C. The following antibodies were used for ChIP: anti-CTCF (Millipore, Cat# 07-729), anti-H3K27ac (Millipore, Cat# 07-449), anti-YY1 (Abcam, Cat# ab109237), anti-RNAPolII (Active Motif, Cat# 39097). 10 µg of antibody was used for each 8-10 million cell pellets.

Beads were washed once in IP wash buffer 1 (20 mM Tris pH 8.0, 2 mM EDTA, 50 mM NaCl, 1 % (v/v) TritonX-100, 0.1 % (w/v) SDS), twice in high-salt buffer (20 mM Tris pH 8.0, 2 mM EDTA, 500 mM NaCl, 1 % (v/v) TritonX-100, 0.01 % (w/v) SDS), once in IP wash buffer 2 (10 mM Tris pH 8.0, 1 mM EDTA, 250 mM lithium chloride, 1 % (v/v) NP-40, 1 % (w/v/) sodium deoxycholate), and twice in 1 x TE. All wash buffers were pre-chilled to ~4 °C. Chromatin was eluted from beads in elution buffer (1 % (w/v) SDS, 100 mM sodium bicarbonate), followed by RNA degradation by adding RNaseA (Roche, #10109169001) at 65°C for 1 hr. Residual proteins were removed and reverse crosslinking of DNA was induced by adding 2.4 U of Proteinase K (NEB, #P8107S) to eluent and incubation at 65°C overnight. The ChIP DNA was purified and extracted using conventional phenol:chloroform extraction and ethanol precipitation methods. ChIP DNA was stored at -20 °C until library preparation. 5 ng of purified ChIP-seq DNA was used for downstream library preparation as described under the "Library preparation (ChIP-seq, Hi-C)".

***Hi-C***

Hi-C was performed on ~2 million cross-linked cells per replicate per condition using the Arima Hi-C kit (Arima Genomics, Inc., #A510008) according to the manufacturer's recommended protocol. Following restriction digest of the chromatin with multiple enzymes, the 5'-overhangs were filled in to label the digested ends with a biotinylated nucleotide. Spatially proximal digested ends of DNA were ligated and then proximity-ligated DNA were purified and sheared to an average size of ~400 bp using a Covaris S220 sonicator at 140 W peak incident power, 10% duty factor, and 200 cycles per burst for 55 seconds. Size-selection and bead purification was performed to obtain 200-600 bp DNA fragments using DNA Purification Beads (AMPure XP Beads, Beckman Coulter, #A63881). Biotin-tagged ligation junctions were enriched after size-selection using Enrichment Beads provided in Arima-Hi-C kit. Streptavidin beads containing enriched DNA fragments were kept up to 3 days at -20 °C, before proceeding with library preparation as described under the "Library preparation (ChIP-seq, Hi-C)".

***Library preparation (ChIP-seq, Hi-C)***

Sequencing libraries were prepared using the NEB Next Ultra II Library Prep Kit (NEB, #E7645S) following the manufacturer's protocol, with minor modifications. For ChIP and Hi-C experiments, end-repair and dA-tailing of DNA was carried out according to the manufacturer's protocol. For Hi-C libraries, adaptor-ligated Hi-C libraries were washed on streptavidin beads twice in 150 µl of wash buffer at 55 °C and once in 100 µl of elution buffer (Arima Genomics, #A510008). Ligation products were eluted from streptavidin beads by boiling at 98°C for 10 min in 15 µl elution buffer. For ChIP-seq libraries, size-selection of adaptor-ligated libraries were performed using AgenCourt Ampure XP beads (Beckman Coulter, #A63881). Library amplification was carried out using NEBNext Ultra II DNA Library Prep Kit for Illumina (NEB, #E7645S) according to the manufacturer's protocol. For ChIP-seq, DNA fragments <1 kb size were size-selected and amplified using 7-8 PCR cycles. After additional purification using AgenCourt Ampure XP beads (Beckman Coulter, #A63881), we assessed the quality of the individual libraries using Agilent Bioanalyzer High Sensitivity DNA Analysis Kits (Agilent, #5067-4626) and quantified DNA concentration using a Kapa Library Quantification Kit (KAPA Biosytems, #KK4835). Libraries were pooled and sequenced on an Illumina NextSeq 500 instrument using 75-cycle v2.5 High Output kit (Illumina, Cat# 20024906) for 75 bp single-end reads for ChIP-seq or 37 bp pair-end for Hi-C.

***hg38 RefSeq Reference transcriptome***

The hg38 reference transcriptome was downloaded on July 7$^{th}$, 2021 using UCSC Table Browser with NCBI RefSeq and RefSeq Curated as track and table options respectively (https://genome.ucsc.edu/cgibin/hgTables?hgsid=1345872709_tycZ8naeqyTXL51A2BV9FK8CsBk0). Genes that (i) are in canonical chromosomes 1-22 and chromosome X, (ii) are not present in multiple chromosomes, (iii) have a gene length greater than 10 kb (iv) are not intersecting the GRCh38 blacklist (https://www.encodeproject.org/files/ENCFF356LFX/), centromeres, or telomeres were analyzed.

***RNA-seq Analysis***

RNA-seq paired-end reads to the above described hg38 RefSeq reference transcriptome were aligned using kallisto (v0.46.1) quant with 100 bootstraps of gene quantification[8]. Following pseudoalignment by kallisto, quantifications of estimated counts were converted into DESeq2 format in R using the library ("tximportData") according to DESeq2 documentation recommendations[7]. Normalized counts were computed for each replicate and each cellular stage using DESeq2 (v1.22.1). Genes with the same transcriptional start sites (TSSs) and transcriptional end sites (TESs) were merged into transcriptional units and the counts were summed together. The final reference transcriptome had 27,030 transcriptional units.

***Hi-C Pre-Processing***

37 bp paired-end reads were aligned to the hg38 genome using bowtie2. Through the HiC-Pro software, the following default global parameters were used: --very-sensitive –L 30 --score-min L,-0.6,-0.2 –end-to-end–reorder and the following local parameters:–very-sensitive –L 20 –scoremin L,-0.6,-0.2 –end-to-end–reorder. unmapped reads, non-uniquely mapped reads, and PCR duplicates were filtered. Cis-contact matrices were assembled by binning paired reads into uniform 10 kb bins using custom scripts and then merged across replicates.

To normalize for sequencing depth, samples were normalized using size factors conditioned on genomic distance from the diagonal. A size factor was computed for each sample by summing the counts of bin-bin pairs that are the same genomic distance and then dividing by the geometric mean. Then, the normalized matrices were rounded to the nearest whole number for downstream loop calling. Poorly mapped regions were removed from the normalized matrices based on an hg38 36-mer alignability track generated using the GEM-mappability alignment software. Bins were set to NaN if the average mappability of a 50kb window centered on that bin was below 50%. In addition, a high outlier filter was implemented to further prevent balancing artifacts. Pixels in the raw contact matrices that exhibited high fold changes relative to a neighborhood of adjacent pixels after balancing were removed. The neighborhood around a given pixel defined by a 5 x 5 square footprint was used to determine a local median based on the values nearby pixels. If the value of a given pixel was greater than 4 times the local median or greater than 4 if the local median was less than 1, then that pixel was determined to be an outlier. Rows containing less than 35 non-zero pixels within 750 kb of the diagonal were removed from further Hi-C analysis. After filtering, matrices were balanced using the Knight-Ruiz matrix balancing algorithm and the final bias factors were retained for downstream loop calling.

***Hi-C Loop Calling: Expected Modeling***

An expected modeling strategy based on the HiCCUPS approach from our previous work and work of others was implemented[12,17,19,22-25]. For expected modeling and all subsequent loop calling steps, the analysis was restricted to bin-bin pairs with interaction distances within 10 Mb. Expected modeling computations were performed on Knight-Ruiz balanced 10kb contact matrices.

First, to account for the distance dependence of Hi-C signal, a one-dimensional expected model, $D$, was computed by averaging interaction counts for each of the first 1000 diagonals spanning 10 Mb (**Equation 1**):

$$D_d = \left(S_{a,b}\right) \ \forall \ d \ such \ that \ 0 \leq d \leq 1000 \quad \textbf{(1)}$$

where $D_d$ is the expected value for interactions between bin-bin pairs $(a, b)$ separated by $d$ bins and $S$ is the balanced contact matrix with a pseudocount of 1 added to each count. Then to correct the one-dimensional expected model for local background signals, each expected value $D_{i,j}$ was multiplied by five separate correction factors (**Equations 2, 4, 6, 8, 10**). These correction factors were computed by summing all bin-bin pairs $(a, b)$ that fall within a geometric footprint centered on pixel $i, j$ in the balanced matrix $S$ and the one-dimensional expected matrix $D$ then finding the ratio between the two sums. The five geometric footprints are the donut footprint (**Equations 2-3**),

$$E_{i,j}^{DF} = D_{i,j} \times \frac{\sum_{(a,b)\in DF_{i,j}} S_{a,b}}{\sum_{(a,b)\in DF_{i,j}} D_{b-a}} \quad \textbf{(2)}$$

$$DF_{i,j} = \{(a,b) \mid (|a-i| \le w) \wedge (|b-i| \le w) \wedge \ (a \ne i) \wedge (b \ne j) \wedge ((|a-i| > p) \vee (|b-j| > p))\}$$

**(3)**

the lower left footprint (**Equations 4-5),**

$$E_{i,j}^{LLF} = D_{i,j} \times \frac{\sum_{(a,b)\in LLF_{i,j}} S_{a,b}}{\sum_{(a,b)\in LLF_{i,j}} D_{b-a}} \quad \textbf{(4)}$$

$$LLF_{i,j} = \{ (a,b) \in DF_{i,j} \mid (a < i) \wedge (b < j) \} \quad \textbf{(5)}$$

vertical footprint (**Equations 6-7**),

$$E_{i,j}^{VF} = D_{I,j} \times \frac{\sum_{(a,b)\in VF_{i,j}} S_{a,b}}{\sum_{(a,b)\in VF_{i,j}} D_{b-a}} \quad \textbf{(6)}$$

$$VF_{i,j} = \{(a,b) | ((b = j-1) \vee (b = j) \vee (b = j+1)) \wedge (|a-i| > p) \wedge (|a-i| \le w)\} \quad \textbf{(7)}$$

horizontal footprint (**Equations 8-9**),

$$E_{i,j}^{HF} = D_{i,j} \times \frac{\sum_{(a,b)\in HF_{i,j}} S_{a,b}}{\sum_{(a,b)\in HF_{i,j}} D_{b-a}} \quad \textbf{(8)}$$

$$HF_{i,j} = \{(a,b) \mid ((a = i-1) \vee (a = i) \vee (a = i+1)) \wedge (|b-j| > p) \wedge (|b-j| \le w)\} \quad \textbf{(9)}$$

and the upper triangle footprint (**Equations 10-17**),

$$E_{i,j}^{UTF} = D_{i,j} \times \frac{\sum_{(a,b)\in UTF_{i,j}} S_{a,b}}{\sum_{(a,b)\in UTF_{i,j}} D_{b-a}} \quad \textbf{(10)}$$

$$UTF_{i,j} = \{ (a,b) \in DF_{i,j} \mid b - a \ge j - i \} \quad \textbf{(11)}$$

respectively. All geometric footprints were parameterized by p=4 and w=10. An additional parameter was implemented where the minimum fraction of finite counts within a footprint must be greater than 0.2 for an expected value to be computed. The final expected value of pixel $i, j$ was computed by finding $E_{i,j}^{UTF}$ when the interaction distance between bins was less than 400kb or finding the maximum expected value across the donut, lower left, vertical, and horizontal footprints when the interaction distance between bins was greater than 400kb but less than 10 Mb (**Equation 12**).

$$E_{i,j} = \begin{cases} E_{i,j}^{UTF}, & for\ b - a \leq 40 \\ max\left(E_{i,j}^{DF}, E_{i,j}^{LLF}, E_{i,j}^{HF}, E_{i,j}^{VF}\right), & for\ 40 < b - a < 1000 \end{cases} \quad \textbf{(12)}$$

**Hi-C Loop Calling: P-values**

The final expected value $E_{i,j}$ and the bias vector $c$ from Knight-Ruiz balancing was used to compute an integer biased expected value for comparison with the integer sequencing depth normalized count $X_{i,j}$ (**Equation 13**):

$$E_{i,j}^{biased} = E_{i,j} \times c_i \times c_j \quad \textbf{(13)}$$

A p-value $P_{i,j}$ was computed by testing the null hypothesis that $X_{i,j}$ was less than or equal to a Poisson-distributed random variable $X_{i,j}'$ with mean $E_{i,j}^{biased}$ (**Equation 14**):

$$P_{i,j} = P\left(X_{i,j} \leq X_{i,j}'\right); \quad X_{i,j}' \sim Poisson\left(E_{i,j}^{biased}\right) \quad \textbf{(14)}$$

**Hi-C Loop Calling: Multiple testing correction**

The lambda-chunking strategy from Aiden[26] was applied for multiple testing correction. First, we stratified bin-bin pairs $(i, j)$ according to their biased expected values $E_{i,j}^{biased}$ using logarithmically spaced bins with a bin spacing $2^{1/3}$. This was followed by a Benjamini-Hochberg false discovery rate control for each chunk separately to obtain a q-value, $Q_{i,j}$, which represent the maximum false discovery rate (FDR) at which an interaction would be called significant.

**Hi-C Loop Calling: Clustering**

After computing $Q_{i,j}$, clusters of nearby significant bin-bin pairs were identified. First, an initial set of significant bin-bin pairs was identified using: (1) a q-value $Q_{i,j} \leq 0.05$ (false discovery rate of 5%), a balanced contact value $S_{i,j} \geq 6$ and an observed over expected fold-change $FC \geq 1.5$. To further reduce the possibility of false positives, clusters with fewer than three significant bin-bin pairs were removed. Large "superclusters", composed of smaller clusters that are more likely to represent individual looping interactions, were found in the initial calls. Therefore, superclusters were split by applying a progressively more stringent q-value threshold on the order of magnitude from 0.05 to 5e-6 FDR. If a cluster became smaller when the threshold was tightened, the smaller, more refined cluster was kept. If a cluster was lost entirely when the threshold was tightened, the cluster right before it was lost was kept. Finally, to further reduce the possibility of false positive interactions being called near the diagonal of the contact matrix, all refined clusters containing a bin-bin pair whose interaction distance was within 3 bins of diagonal were removed.

**Hi-C Differential Loop Calling**

To call differential loops between conditions, 3DeFDR-HiC was used[12]. In brief, 3DeFDR-HiC performs a negative binomial likelihood ratio test for every pixel engaged in loops genome-wide. The negative binomial model was parameterized by (i) the mean count per pixel across replicates for the two cell stages being compared (ii) a distance-dependent scaling factor and (iii) the Knight-Ruiz balancing bias factor, and (iv) an estimated distance dispersion per pixel across replicates for the two cell stages being compared. Then a likelihood ratio test was performed to obtain p-values against the null hypothesis that each looping pixel was not differential. The Benjamini-Hochberg procedure was applied to correct these p-values for multiple testing using an FDR threshold of 30%. Once differential pixels were identified, they were re-clustered back into their original clusters and compared for occupancy. If a cluster had more than 75% of its pixels as being condition specific, then that cluster was classified as condition specific. Otherwise, the cluster was classified as invariant. To visually confirm differential loop calls, aggregate peak analysis was performed based on the mean Hi-C signal of a +/- 130 kb region centered on the differential loop calls.

**ChIP-seq Analysis**

RNA Polymerase II (RNAPolII), CTCF, and H3K27ac ChIP-seq reads were aligned to the hg38 genome using Bowtie. Reads with more than two possible alignments were removed. To account for sequencing depth differences, all immunoprecipitation libraries were downsampled to 24 million reads. Input libraries were also downsampled to 24 million reads. MACS2 callpeak was used to call peaks with a p-value cutoff of $1 \times 10^{-4}$ for RNAPolII, CTCF, and H3K27ac, and a broad peak cutoff of $1 \times 10^{-4}$ for RNAPolII and H3K27ac. Bedgraph pileups were also generated using the -B and --SPMR flags and then converted to bigwigs using UCSC bedGraphToBigWig. During preliminary analysis, it was noted the RNAPolII, CTCF, and H3K27ac ChIP-seq signals were not proportional across libraries. To address this, each pileup interval was divided by scalar size factors. Each size factor was determined by finding the maximum global signal for each chromosome, finding the mean of those maximums, and then dividing that mean by 14.106 for the RNAPolII libraries, 11.339 for the CTCF libraries, and 9.965 for the H3K27ac libraries.

**RNAPolII Occupancy Analysis**

RNAPolII binding was profiled for each gene by computing the transcription start site (TSS) ChIP-seq pileup signal ($S_{TSS}$), the transcription end site (TES) ChIP-seq pileup signal ($S_{TES}$), and the gene body ChIP-seq pileup signal ($S_{GB}$). The total pileup signal spanning +/- 500 bins relative to the TSS was obtained using the values function from the pyBigWig python package (v0.3.18). Then, $S_{TSS}$ was computed as the maximum ChIP-seq pileup signal within this region $S_i$ (**Equation 15**):

$$S_{TSS} = \max_{i \in \{-500\ bp \ldots 500\ bp\}} (S_i) \tag{15}$$

To correct for enriched 3' signal, the total pileup signal spanning +/- (gene length * 0.025) relative to the TES using the pyBigWig values function was also obtained. $S_{TES}$ was computed as the mean ChIP-seq pileup signal within this region $S_j$ (**Equation 16**):

$$S_{TES} = \underset{j \in \{-(0.025 * gene\ length)\ bp \ldots +(0.025 * gene\ length)\ bp\}}{mean} (S_j) \tag{16}$$

Finally, the total pileup signal spanning +500 base-pairs from the TSS region to the start of the TES region was obtained. Before computing the gene body pileup signal, $S_{GB}$, the region was resized into 4,375 bins to normalize for gene length. This was accomplished using the cv2.resize function from the opencv-python package (v4.5.4.58). After resizing, the gene body signal, $S_{GB}$, was computed (**Equation 17**):

$$S_{GB} = \underset{k\,\in\{0\ bins \dots 4375\ bins\}}{mean}(S_k) \tag{17}$$

Each gene was parsed into repressed, elongated, and initiated categories across each cellular stage. For a gene to be classified as repressed, four conditions must be met: $S_{TSS} < 0.75$ and $S_{GB} < 0.2$, $S_{TES} < 0.3$ and mean normalized gene expression less than 300. For a gene to be classified as initiated, three conditions must be met: $S_{TSS} > 0.75$ and $S_{GB} < 0.2$, and $S_{TES} < 0.3$. For a gene to be classified as elongated, two conditions must be met: $S_{TSS} > 0.75$ and $S_{GB} > 0.25$. ChIP-seq pileup heatmaps were generated using deeptools.

**Intersection of RNAPolII Occupancy Profiles with CTCF peaks**

For each gene that exhibits each of the three RNAPolII occupancy profiles in each cellular stage, promoter regions (i.e. 2kb upstream of the TSS) were intersected with CTCF peaks identified in that cellular stage using bedtools intersect. For example, the promoters of genes that were classified as elongated in NPCs were intersected with CTCF peaks identified in NPCs, the promoters of genes that were classified as elongated in neuron were intersected with CTCF peaks identified in neuron, et cetera.

**Gene Ontology Analysis**

Gene ontology enrichment was performed using WebGestalt (http://www.webgestalt.org). The following settings were used: Organism of interest = Homo Sapiens; Method of interest = Over-Representation Analysis; Functional Database = geneontology, Biological Process noRedundant; Select Reference Set, genome.

**Parsing RNAPolII Transition Classes**

For each cellular stage transition (i.e. hiPSC-to-NPC and NPC-to-neuron), each gene was parsed into nine RNAPolII transition classes based on their RNAPolII occupancy profiles. Genes were parsed as follows:

i. **Repressed-Repressed:** RNAPolII occupancy profile is classified as repressed in both prior and latter cellular stages.

ii. **Repressed-Initiated**: RNAPolII occupancy profile is classified as repressed in the prior cellular stage but initiated in the latter cellular stage.

iii. **Repressed-Elongated:** RNAPolII occupancy profile is classified as repressed in the prior cellular stage but elongated in the latter cellular stage.

iv. **Initiated-Initiated**: RNAPolII occupancy profile is classified as initiated in both prior and latter cellular stages.

v. **Initiated-Repressed**: RNAPolII occupancy profile is classified as initiated in the prior cellular stage but repressed in the latter cellular stage.

vi. **Initiated-Elongated**: RNAPolII occupancy profile is classified as initiated in the prior cellular stage but elongated in the latter cellular stage.

vii. **Elongated-Elongated**: RNAPolII occupancy profile is classified as elongated in both prior and latter cellular stages.

viii. **Elongated-Initiated**: RNAPolII occupancy profile is classified as elongated in the prior cellular stage but initiated in the latter cellular stage.

ix. **Elongated-Repressed**: RNAPolII occupancy profile is classified as elongated in the prior cellular stage but repressed in the latter cellular stage.

**Parsing Looping Classes**

For each RNAPolII transition class, promoters (i.e. 2kb upstream of the TSS) were intersected with both anchors of each differential and invariant loop calls (Figure 4, Supplemental Figure 3).

For the hiPSC-to-NPC transition, a gene was classified as **Class 1** if the promoter intersected with one or more hiPSC-specific loops, **Class 2** if the promoter only intersected with one or more NPC-specific loops, **Class 3** if the promoter intersected with one or more hiPSC-specific loops and one or more NPC-specific loops, **Class 4** if the promoter intersected with only one or more invariant loops, and **Class 5** if the promoter did not intersect with any differential or invariant loops.

For the NPC-to-neuron transition, a gene was classified as **Class 1** if the promoter intersected with one or more NPC-specific loops, **Class 2** if the promoter only intersected with one or more neuron-specific loops, **Class 3** if the promoter intersected with one or more NPC-specific loops and one or more neuron-specific loops, **Class 4** if the promoter intersected with only one or more invariant loops , and **Class 5** if the promoter did not intersect with any differential or invariant loops.

**Defining Differential Enhancer Regions**

For Figure 5 and Supplemental Figure 7, we filtered genes parsed by RNAPolII and looping status based on their proximity to differential enhancers. To identify differential enhancers, concatenated list of H3K27ac ChIP-seq peak calls identified in NPC and neurons were created. From this list, any H3K27ac peaks that overlapped hg38 TSS±1kb, hg38 Refseq exons, 3' UTRs and 5' UTRs (downloaded from the UCSC Genome browser on October 25$^{th}$ 2022) were removed using bedtools. This resulted in H3K27ac peaks at only introns and intergenic regions. Finally, H3K27ac peaks were parsed into cell-type specific enhancers and invariant enhancers in the following way:(i) calculating the average bigwig signal across the peak interval using the pybigwig package (v0.3.13) in both conditions, and (ii) calculating the fold change in H3K27ac-seq signal as [NPC stage/neuron stage]. A peak was assigned as an NPC-specific enhancer if it exhibited a fold change H3K27ac ChIP-seq signal as [NPC stage/neuron stage] > 2.0 and if the NPC signal at the peak was above 20th percentile signal threshold of all the enhancers. A peak was assigned as a neuron-specific enhancer peak in the same manner with the conditions reversed. The remaining enhancer peaks were classified as invariant enhancers (i.e the peaks with less than two-fold-change in signal between the two conditions and if the signal under the peak was above 20th percentile signal threshold of all peaks in both conditions).

The same method was applied to the H3K27ac ChIP-seq peak calls identified in hiPSC and NPCs. A peak was assigned as an hiPSC-specific enhancer if it exhibited a fold change H3K27ac ChIP-seq signal as [hiPSC stage/ NPC stage] > 2.0 and if the hiPSC signal at the peak was above 20th percentile signal threshold of all the enhancers. A peak was assigned as an NPC-specific enhancer peak in the same manner with the conditions reversed.

**Classification of NPC-specific and neuron-specific loops using promoter positions and differential H3K27ac enhancer regions**

Loops formed by the genes present in the RNAPolII transition classes based on the cell-type specific H3K27ac enhancers from ChIP-seq peak calls were classified (see previous section). Loop anchors were defined as the region between start and end coordinates of the two sides of the rectangle enclosing all the bin-bin pairs in the looping cluster. Promoter regions were defined as 2kb upstream of the TSS of genes.

Promoter loops were classified as follows:

i. **Promoter–neuron-specific enhancer**: if one anchor overlapped a promoter region and the other anchor overlapped a neuron-specific enhancer regardless of the presence of other enhancers.

ii. **Promoter–NPC-specific enhancer**: if one anchor overlapped a promoter region and the other anchor overlapped a prior NPC-specific enhancer but not a neuron-specific enhancer.

iii. **Promoter–Cell type-invariant enhancer**: if one anchor overlapped a promoter region and the other anchor overlapped only invariant enhancers.

iv. **Promoter–Non-coding // non-enhancer**: if one anchor overlapped a promoter region and the other anchor did not overlap with any enhancers.

v. **Promoter–Promoter:** if both anchors were overlapping any promoter regions as defined above. This group is also mutually exclusive with the promoter-enhancer groups.

For the hiPSC-to-NPC transition, we classified promoter loops as follows:

i. **Promoter–NPC-specific enhancer**: if one anchor overlapped a promoter region and the other anchor overlapped an NPC-specific enhancer regardless of the presence of other enhancers.

ii. **Promoter–iPSC-specific enhancer**: if one anchor overlapped a promoter region and the other anchor overlapped a prior hiPSC-specific enhancer but not a NPC-specific enhancer.

iii. **Promoter–Cell type-invariant enhancer**: if one anchor overlapped a promoter region and the other anchor overlapped only invariant enhancers.

iv. **Promoter–Non-coding // non-enhancer**: if one anchor overlapped a promoter region and the other anchor did not overlap with any enhancers.

v. **Promoter–Promoter:** if both anchors were overlapping any promoter regions as defined above. This group is also mutually exclusive with the promoter-enhancer groups.

**Stratification of promoter-promoter looping genes and non-looping genes**

Promoter-promoter looping genes identified in the previous section and non-looping genes identified in the "Parsing Looping Classes" section were further stratified based on the presence of close-range intronic and intergenic H3K27ac peaks (i.e. within ±80 kb of the TSS). This range was determined based on the resolution of the loop calling method. Genes that did not have any intronic and intergenic H3K72ac peaks within ±80 kb of the TSS were further characterized for gene expression in Figure 5 and Supplemental Figure 7.

**RNAPolII Occupancy Analysis of DLD-1 RNAPolII ChIP-seq**

First, the RNAPolII ChIP-seq bigwig from[27] was lifted over from hg19 to hg38 using UCSC liftOver. Then, from the hg38 lifted bigwig, genes were classified as repressed, initiated and elongated using the algorithm described in the RNAPolII Occupancy Analysis section. For a gene to be classified as repressed, three conditions must be met: $S_{TSS} < 0.75$ and $S_{GB} < 0.1$, $S_{TES} < 0.3$. For a gene to be classified as initiated, three conditions must be met: $S_{TSS} > 0.75$ and $S_{GB} < 0.1$, and $S_{TES} < 0.3$. For a gene to be classified as elongated, two conditions must be met: $S_{TSS} > 0.75$ and $S_{GB} > 0.2$.

**Analysis of DLD-1 CTCF CUT&Tag**

MACS2 bdgpeakcall was used to call CTCF peaks from the published bedgraphs of replicates 1 and 2 separately[28]. A p-value cutoff of $1 \times 10^{-2}$ was used. Peak calls were then merged using bedtools merge. For each gene classified as repressed, initiated, or elongated, the promoter regions (i.e. 2kb upstream of the TSS) were intersected with the merged CTCF peaks using bedtools intersect.

**Analysis of DLD-1 Micro-C**

Using the provided mcool files from[28], the Mustache package was used to identify loops in Micro-C from control DLD-1 cells without RNAPolII depletion. The following parameters were used: -pt 0.05 and -r 5000. The loop epicenters that intersected the 2kb promoters of repressed, initiated, and elongated genes with and without CTCF were used in the paired aggregate peak analyses in Figure 7. ChIP-seq pileup heatmaps of gene promoters that intersected loops called in the control condition in Figure 7 were generated using deeptools.
